# Metabolic trade-offs in sugar beet under drought and beet leaf miner infestation: implications for herbivore success

**DOI:** 10.64898/2026.03.01.708914

**Authors:** Shahinoor Rahman, Musrat Zahan Surovy, Ilka Vosteen, Michael Rostás

## Abstract

Increasing frequency of drought under climate change threatens crop production and intensifies pest pressures, yet the interactive effects of drought and herbivory on plant metabolism and ecological outcomes remain incompletely understood. We subjected sugar beet (*Beta vulgaris*) plants to moderate and high drought, alone or with infestation by the beet leaf miner (*Pegomya cunicularia*), and analyzed plant physiology, central metabolites, and volatile organic compound (VOC) emissions. Drought alone reduced growth and photosynthetic efficiency, while combined stress led to accentuated metabolic reprogramming, including increased amino acids and organic acids, and a concurrent suppression and alteration of VOC emissions, especially in plants affected by high drought and leaf mining. The resulting changes in VOC blends reduced plant attractiveness to ovipositing females, leading to fewer eggs laid on severely stressed plants. Contrastingly, moderate drought generated a nutrient-rich environment: larvae feeding on these plants exhibited the highest growth rates, larger pupae and adults, and increased feeding damage. High drought strongly limited both plant water content and larval development. These findings reveal a stress-dependent tradeoff between enhanced leaf nutritional quality and reduced host detectability, underscoring the importance of integrating multi-stress plant biology for future pest management and crop resilience.

**Highlight:** Combined drought and herbivory in sugar beet plant triggered stress-intensity-dependent trade-offs between leaf nutritional quality and volatile emissions, affecting beet leaf miner performance and oviposition—highlighting how multi-stress interactions shape plant–insect dynamics.

Graphical Abstract

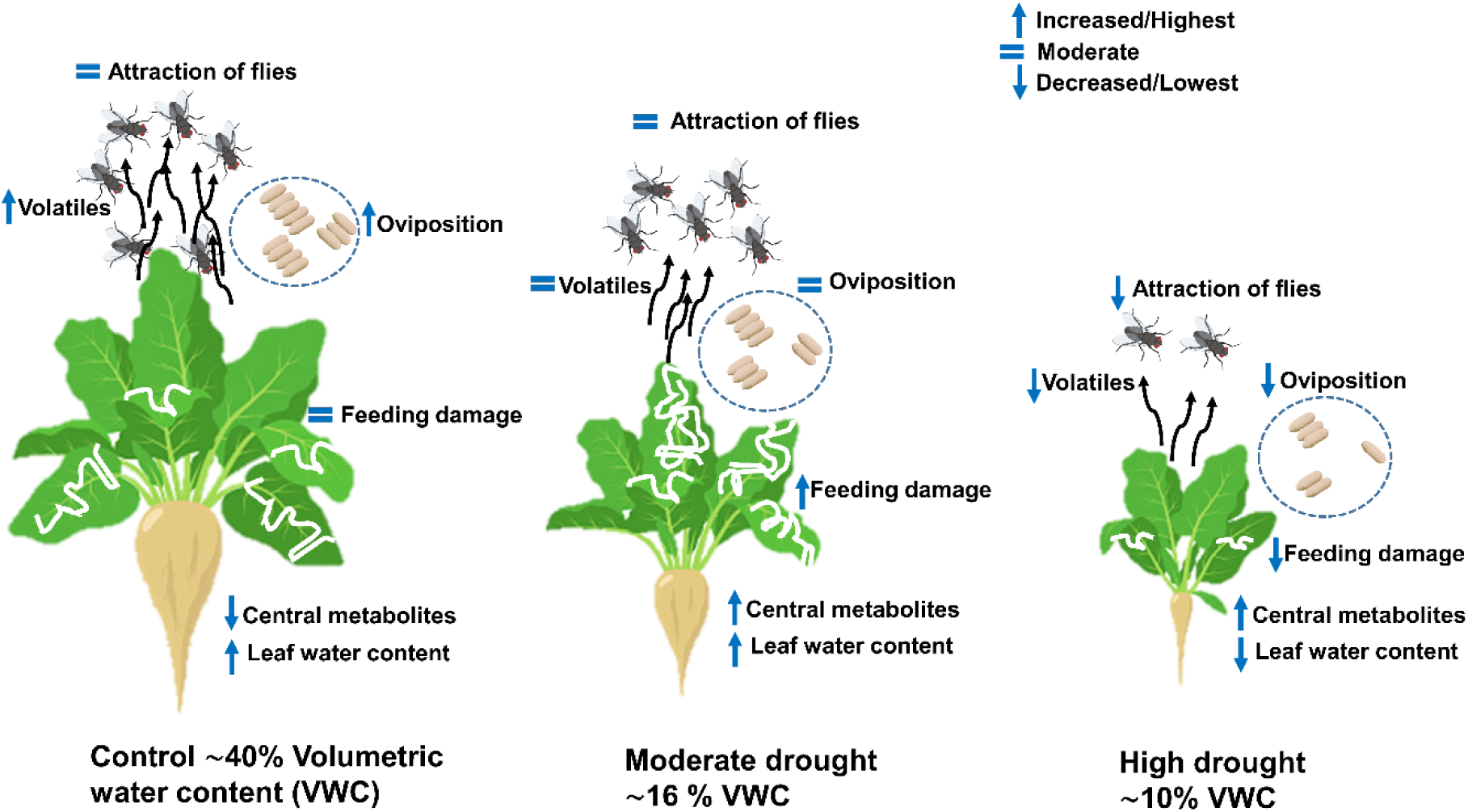

## Introduction

Drought is a major abiotic stressor limiting sustainable crop production (Ray *et al*., 2020), with climate change driving more frequent and severe drought episodes globally, as recently experienced in central Europe (Boergens *et al*., 2020; Haslinger and Mayer, 2023). These water deficits profoundly impact plant physiology, triggering complex adjustments in growth, metabolism, and defense (Bhattacharya 2021). Importantly, drought rarely occurs in isolation; crops are often simultaneously challenged by biotic stressors such as insect herbivores. The interactive effects of combined abiotic and biotic stresses can fundamentally reshape plant responses and influence plant–insect ecological dynamics (Hamann *et al*., 2021; Lin *et al*., 2023), yet remain insufficiently understood at the mechanistic level.

Drought stress fundamentally limits plant growth and development by reducing plant height, biomass, leaf area, and water content, while inducing oxidative stress that impairs photosynthetic efficiency (Yao et *al.*, 2018; Zhuang *et al*., 2020; Yang *et al*., 2021). In response, plants activate a suite of physiological and molecular mechanisms—including the accumulation of osmoprotectants such as soluble sugars and proline—to maintain cellular homeostasis under water deficit (Anjum *et al*., 2017; Sharma *et al*., 2019). These adjustments drive broad metabolic reprogramming, shifting the balance of primary metabolites and potentially diverting resources away from specialized defense pathways (Wittstock and Gershenzon, 2002; Fàbregas and Fernie, 2019). Such metabolic changes not only affect plant vigor but also alter tissue nutritional quality, which can influence herbivore growth and performance (Mevi-Schütz *et. al.*, 2003, Boggs and Freeman, 2005). However, the effects of drought on plant–insect interactions are highly context-dependent and can either enhance or suppress herbivore success, depending on the severity of stress, insect feeding strategy, and plant–insect specificity (Kuczyk *et al*., 2021; Shehzad *et al*., 2021; Carvajal Acosta *et al*., 2023). When drought occurs in combination with biotic stressors such as herbivory, plant resource allocation and defense strategies may be further modified, with consequences for both plant resilience and insect performance.

Drought-induced oxidative stress impacts photosynthetic efficiency, restricting carbon assimilation and thus the carbon pool available for the biosynthesis of both central and specialized metabolites, including defense compounds and volatile organic compounds (VOCs). The intricate biosynthesis of secondary metabolites like VOCs is often tightly linked to central metabolic pathways (Pott *et al*. 2019). Under carbon-limiting conditions caused by drought, plants may strategically reallocate resources, potentially altering VOC profiles, leading to either increased or decreased production of specific compounds depending on the plant species and stress intensity (Szabó *et al*., 2020). The biosynthesis and emission of VOCs are not governed solely by metabolic fluxes but are also strongly influenced by physiological parameters such as stomatal conductance and leaf water status. The emission rate of compounds with low Henry’s law volatility constant (H*^pc^*, Sandra 2015), such as alcohols, carbonyls, and oxygenated monoterpenes, can be particularly sensitive to stomatal closure, which often accompanies drought stress (Lin *et al*., 2022). As a result, drought can alter not only the total amount of VOCs released but also the qualitative composition of the volatile blend, with some compounds being upregulated and others downregulated (Tariq *et al*., 2013; Weldegergis *et al*., 2015). These drought-induced changes in VOC profiles are ecologically significant, as herbivorous insects rely on plant volatiles as cues for host location (Bruce *et al*., 2005; Bruce and Pickett, 2011), and females often use these signals to select suitable oviposition sites (Honda, 1995). Modifying the relative proportions of components within a VOC blend can alter the blend’s informational content for insects (Rusman *et al*. 2024). Furthermore, plant VOC emission is not only constitutive but also inducible by biotic stressors such as herbivory (Bezerra *et al*., 2021), leading to complex and dynamic changes in volatile profiles during simultaneous exposure to both drought and insect attack. The interplay of drought and herbivory can generate distinct VOC profiles that alter herbivore attraction. Thus, deciphering how combined stresses shape plant volatiles is essential for predicting plant–insect interactions and informing pest management under climate change.

Sugar beet (*Beta vulgaris*) is a major temperate crop whose productivity is increasingly threatened by both climatic and regulatory changes. The recent rise in drought frequency, coupled with restrictions on chemical pest control such as neonicotinoid bans, has intensified pest pressures in sugar beet cultivation (Ebmeyer *et al*., 2021; Viric Gasparic *et al*., 2021). The beet leaf miner (*Pegomya cunicularia*) is a significant pest whose larvae create large, irregular blotch-shaped mines in sugar beet leaves, sometimes causing substantial damage (Michelsen, 1980; Hilman, 2022). Drought-induced changes in plant nutritional status and secondary metabolite profiles may alter the suitability of sugar beet as a host, with potential consequences for herbivore development and reproduction. Despite this, the mechanistic basis of how combined drought and herbivory shape sugar beet physiology, metabolism, and volatile emissions—and how these plant responses mediate interactions with herbivores—remains poorly understood. Despite growing interest in drought–herbivory interactions, most studies have treated drought as a binary factor (stress vs. no stress), limiting our understanding of how increasing drought intensity shapes plant defense strategies. In particular, it remains unclear whether moderate and severe drought differentially alter the coordination between primary metabolism, volatile-mediated indirect defenses, and herbivore performance. Furthermore, relatively few studies have integrated central metabolomic profiling, volatile emissions, herbivore development, and behavioural responses of both herbivores and their natural enemies within a single experimental framework. This limits our mechanistic understanding of how plants prioritize resource allocation and defense coordination under combined abiotic and biotic stress.

In this study, we addressed this knowledge gap using an integrative experimental framework that combines central metabolomics, volatilomics, and insect behavioral and performance assays to disentangle the individual and interactive effects of two drought intensities and beet leaf miner herbivory on sugar beet. Specifically, we addressed the following questions: (i) How do drought stress and beet leaf miner infestation, individually and in combination, reprogram the central metabolite profiles of sugar beet? (ii) In what ways do these stresses alter the composition and emission of plant volatile organic compounds? and (iii) How do these physiological and chemical changes affect host plant detection, oviposition preference, and subsequent development of *Pegomya cunicularia*? By linking plant metabolic and volatile reprogramming to both herbivore development and oviposition behavior, our study provides novel, mechanistic insight into how simultaneous abiotic and biotic stresses shape plant–insect interactions. By revealing stress-intensity–dependent shifts in plant defense coordination across trophic levels, this study advances mechanistic understanding of plant adaptation to combined environmental stress.

## Materials and Methods

### Insects, plants, drought system, and experimental setup

Adults of the beet leaf miner *Pegomya cunicularia* (Diptera: Anthomyiidae) were collected from sugar beet fields near Göttingen (51°30’19.4”N 9°55’48.3”E), Germany, and mass-reared under laboratory conditions. Detailed rearing protocols are provided in the supporting information file (Supplementary Fig. S1).

Seeds of *B. vulgaris* subspec. *vulgaris* cultivar ‘Vasco’ (SESVanderHave, Belgium) were sown in plastic trays (54 holes, each 3.5 cm diameter) filled with quartz sand (0.2-0.8 mm grain size). All experiments were conducted under controlled climate conditions (16L:8D photoperiod; light intensity: 130 ± 10 µmol m^-2^ s^-1^; relative humidity: 65 ± 5 %; temperature: 20 ± 2 °C). Seedlings were watered with tap water until 11 days after sowing (DAS), then supplied with half-strength modified Hoagland solution (Rahman *et al*. 2025) from 12 to 23 DAS. At 24 DAS, seedlings were transferred to a capillary-based drought system (Marchin *et al*., 2020), which was modified and applied as described by Rahman *et al*., (2025), and drought was implemented at 27 DAS under three regimes: i) control: ∼40% volumetric water content (VWC) ii) moderate drought: ∼16% VWC and iii) high drought ∼10% VWC. Based on preliminary calibration experiments, the maximum water-holding capacity of the quartz sand substrate (0.2–0.8 mm grain size) was determined to be approximately 40% volumetric water content (VWC), which was therefore used as the well-watered control condition. The permanent wilting point of sugar beet in this system occurred below 9% VWC. Moderate (16% VWC) and high (10% VWC) drought treatments were thus selected to impose physiologically relevant but non-lethal water deficits above the wilting threshold.

### Experiment 1: Plant growth and water content

Plant height was measured at three time points: at the two true leaf stage prior to drought treatment (26 days after sowing, DAS), and subsequently at 36 DAS and 46 DAS, across all three treatments (control, moderate drought, and high drought). At 58 DAS plants were harvested and the following parameters were recorded: total biomass, root weight, shoot weight, root length, shoot length, root-to-shoot ratio (by both weight and length), total number of leaves, and average leaf area (measured using an LI-3100C Area Meter, LI-COR Biosciences, Germany).

After measuring the plant morphological parameters, leaf disks (3.5 cm diameter) were excised from two fully expanded leaves per plant. The fresh weight (FW) of the disks was recorded using a high precision balance (KC BA 100, Sartorius Micro, Germany), after which the disks were oven-dried at 60 °C for 72 h (BD-115, Binder, Germany) to obtain their dry weight (DW). Leaf water content was calculated using the following formula:

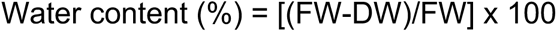

where FW = Fresh weight of the leaf disk, DW = Dry weight of the leaf disk

### Experiment 2: Leaf photosynthetic efficiencies and beet leaf miner development

#### Leaf photosynthetic efficiencies

At 36 DAS, corresponding to 10 days after the initiation of the drought treatments, four freshly laid eggs (≤24 h old) of *P. cunicularia* were carefully placed on the abaxial surface of the second pair of fully expanded leaves using a fine brush, across all three drought conditions (control, moderate drought, and high drought). This full factorial design resulted in six treatment combinations: each drought level with and without beet leaf miner infestation. Larval emergence generally occurred within 5 days following egg placement. Chlorophyll fluorescence parameters were measured at four time points: 39 DAS (pre-infestation), 44 DAS, 49 DAS, and 54 DAS. Prior to each measurement, plants were dark-adapted for 30 minutes. A MINI-PAM-II chlorophyll fluorometer (Heinz Walz GmbH, Germany) was used to assess: maximum photochemical quantum yield of photosystem (PS) II (Fv/Fm), the effective photochemical quantum yield of PS II (Ф*_PSII_*), and electron transport rate (ETR).

#### Beet leaf miner development

Daily monitoring was conducted to record egg hatching, larval emergence, and development time. At 55 DAS, mined leaves were excised, photographed under standardized lighting conditions against a contrasting background, and analyzed using ImageJ to determine mined leaf area. Each image was calibrated using a millimeter reference scale included in the photograph. Total leaf area and mined area were quantified manually using the freehand selection tool. All measurements were conducted individually and by the same operator to ensure consistency. Leaves were then placed in soil-filled rearing boxes (sand:clay:organic matter = 2:1:1) for pupation. Pupal weight was measured using a microbalance (KC BA 100, Sartorius). After adult emergence, flies were anesthetized using CO₂ and weighed. Developmental time from egg to pupa and pupa to adult was recorded.

### Experiment 3: Central metabolome analysis

Leaves were harvested at 42 DAS from plants subjected to drought and leaf miner treatments as described in experiment 2. Larvae were quickly removed, leaves were excised at the base, and immediately flash-frozen in liquid nitrogen. The samples were lyophilized, ground, and stored at −80 °C. For metabolite extraction, 40 mg of powdered leaf tissue was transferred to a 1.5 mL microcentrifuge tube and extracted with 1 mL methanol (LC-MS grade, 99.95 %). The mixture was vortexed (30 s), sonicated (3 min), and centrifuged at 20,156 *g* for 10 min. Supernatants were transferred to glass vials and 20 µL adonitol (20 ng/µL in methanol) was added as internal standard. Samples were dried under vacuum at 30 °C using a rotary vacuum concentrator (RVC 2-25 CD plus) and flushed with argon before overnight storage at −80 °C. For derivatization, 200 µL methoxyamine hydrochloride (20 mg/mL in pyridine) was added to each sample and incubated at room temperature for 90 min. Subsequently, 20 µL MSTFA (N-methyl-N-trimethylsilyl-trifluoroacetamide) was added and vortexed prio to analysis. GC-MS analysis was performed using an Agilent 7890B GC system coupled to a 5977B mass selective detector (MSD), equipped with an Rtx-5 capillary column (30 m × 0.25 mm I.D. × 0.25 µm film thickness). One microliter of the derivatized sample was injected via a PAL RSI 85 autosampler with a split ratio of 20:1. GC oven conditions were as follows: 70 °C for 2 min, ramped to 325 °C, and held for 10 min. Helium was used as the carrier gas at a constant flow rate of 1 mL/min. Retention indices were calculated using mixture of C8–C40 *n*-alkanes. Metabolite identification was performed using MS-DIAL (v4.8) with the Golm Metabolome database (GMD), BinBase, and in-house spectral libraries. Peak areas were normalized to the internal standard (adonitol), which was added after extraction and before derivatization. Thus, normalization accounted for variation arising from derivatization and GC–MS analytical performance. Metabolites that did not match existing database entries were labeled as “unknown”.

### Experiment 4: Analysis of plant volatiles

Volatiles were collected from plants representing the six treatment combinations, using a modified six-arm olfactometer setup (Turlings *et al*., 2004). Plants at 42 DAS, prepared as described in Experiment 2, were enclosed in bags made of plasticizer-free polyester foil (Bratschlauch, Cofresco Frischhalteprodukte GmbH & Co. KG, Germany) for dynamic headspace sampling. VOC collection was conducted over a 24 h period (09:00 to 09:00) under controlled lighting conditions (10 fluorescent PAR lamps; 130 µmol m⁻² s⁻¹ at 3 cm distance) with 16 h light/8 h dark photoperiod. Charcoal-filtered air (two activated charcoal filters, 400 cc, Alltech, Deerfield, IL, USA) from an in-house compressor was humidified (relative humidity ∼56%) by passing through deionized water and supplied to each bag at 1.0 L min⁻¹. Air was simultaneously pulled from the bag at 0.5 L min⁻¹ using a vacuum pump (N816.3KN.18, Laboport®, Germany) through a volatile collection trap (7 cm glass tube) containing 30 mg Porapak Q (80–100 mesh; Volatile Collection Trap LLC, FL, USA). After collection, volatiles were eluted with 150 μL dichloromethane (GC grade) and subjected to GC-MS analysis. Six biological replicates were conducted per treatment group over six consecutive collection days. Dynamic headspace extraction was used to trap plant volatiles, which were subsequently analyzed by GC-MS using an Agilent 7890B gas chromatograph coupled to a 5977B mass selective detector (Agilent Technologies) as described in Rahman *et al*. (2025). Prior to GC–MS injection, 200 ng tetralin (1,2,3,4-tetrahydronaphthalene; Sigma-Aldrich, Taufkirchen, Germany) was added to each sample as an internal standard. A 2 μL aliquot was injected in pulsed splitless mode using an automated injection system. The GC oven temperature was held at 40 °C for 3 min, then increased to 220 °C and held for 10 min. Helium was used as the carrier gas at a constant flow rate of 1.5 mL min⁻¹. Retention indices were calculated using a C8–C20 n-alkane mixture (∼40 mg L⁻¹ each in hexane; Sigma-Aldrich). Tentative compound identification was performed using MSD ChemStation software in combination with the NIST17 and Wiley11 mass spectral libraries, requiring a reverse match score >90%. Experimental retention indices were compared with published values (Van den Dool and Kratz method), and compounds were accepted when RI values matched within ±5 units, and the results are summarized in Supplementary Table S1. Compound quantification was performed by normalizing peak areas to the internal standard (tetralin).

### Experiment 5: Olfactory responses of P. cunicularia flies to sugar beet volatiles

The behavioral responses of adult beet leaf miners to VOCs from treated plants was assessed using a glass Y-tube olfactometer setup. The Y-tube had an internal diameter of 3.5 cm, with each arm measuring 20 cm in length. The setup was housed within a photo light box (60 × 60 × 60 cm, Yorbay eBusiness GmbH, Germany) to avoid visual disturbance of the insects and placed on a green platform inclined at an 8° angle. Illumination was provided by LED lights with a color temperature of 5500 K, creating an even, daylight-like environment.

The plants (42 DAS) used as odor sources were prepared as described in Experiment 2. Before the experiment, they were enclosed in oven bags (Bratschlauch, Toppits, Germany) and connected via 6.4 mm Tygon tubing. Filtered and humidified air was pushed into each odor chamber at 1.0 L min⁻¹ from an in-house compressor. Odor comparisons included: i) Soil substrate vs. control, ii) Moderate drought vs. control, iii) High drought vs. control, iv) Control + beet leaf miner vs. control, v) Moderate drought + beet leaf miner vs. control and vi) High drought + beet leaf miner vs. control. For each odor comparison, naïve female flies (without prior exposure to host plant volatiles) were tested individually (n = 36). Following every 6 flies, the positions of the odor sources were switched to avoid directional bias. After every set of 12 flies, new plant combinations and a clean Y-tube were used. A choice was recorded when a fly entered one of the olfactometer arms within 3 min. All glassware was cleaned with demineralized water and 99.5 % acetone, dried, and heat-sterilized (oven dried) at 180 °C for 2 h before reuse.

### Experiment 6: Oviposition preferences of P. cunicularia

To assess the oviposition preferences of beet leaf miner flies, 42-day-old plants were used in two experimental setups: choice and no-choice. In the choice assays, female flies were presented with multiple treatment options within mesh tents (60 × 60 × 70 cm). Two different configurations were used. In the first, plants from the three drought treatments were placed side by side. In the second, plants representing all six treatment combinations were offered simultaneously: control, control + beet leaf miner, moderate drought, moderate drought + beet leaf miner, high drought, and high drought + beet leaf miner. In the no-choice assays, each tent contained one plant from a single treatment, thereby eliminating any alternative oviposition sites. Plants were not enclosed individually, and females had unrestricted access to plant surfaces, allowing visual, olfactory, and contact cues to contribute to oviposition decisions. In all tents, three naïve 10-day-old *P. cunicularia* females were released and allowed to oviposit freely for 24 h. At the end of the assay period, eggs laid on the plants were counted. Each experimental setup was replicated 18 times.

### Statistical analysis

All data analyses were conducted in R (v4.2.1) using RStudio (v2022.07.02+576) and MetaboAnalyst 5.0 (Pang *et al.,* 2021). Appropriate statistical models were selected based on distribution assumptions, which were validated using residual diagnostics. Details of statistical models, software packages, and test procedures for each experiment are provided in the supporting information file. For significant main effects, post-hoc comparisons were conducted using Tukey’s HSD or Dunn’s test as appropriate.

## Results

### Effects of drought stress on the performance of sugar beet plants

Drought stress significantly reduced sugar beet height over time (GLMM: Drought: χ^2^= 92.19, *p* < 0.001; Time: χ^2^= 281.37, *p* < 0.001). High drought also led to pronounced reductions in total biomass (LMM: F_2,6_= 484.34, *p* < 0.001), root weight (LMM: F_2,6_= 63.50, *p* < 0.001), shoot weight (LMM: F_2,6_= 268.07, *p* < 0.001), and shoot length (Kruskal-Wallis: *χ^2^* _(*df* 2)_= 47.17, *p* < 0.001). However, root length (LMM: F_2,6_= 13.49, *p* = 0.006), root-to-shoot ratio by length (GLMM: χ^2^ _(*df* 2)_= 43.79, *p* < 0.001), and by weight (LMM: F_2,6_= 16.50, *p* = 0.003) was highest in highly-stressed plants. The total number of leaves (Kruskal-Wallis: *χ^2^*_(*df* 2)_= 45.14, *p* < 0.001) and the average leaf area (LMM: F_2,6_= 893.11, *p* < 0.001) were significantly reduced in highly-stressed plants compared to control plants. These plant morphological characteristics were already evident in moderate drought conditions and became more pronounced under high drought (Supplementary Fig. S2 A-J). Moreover, leaf water content significantly decreased under high drought conditions (Kruskal-Wallis: *χ^2^* _(*df* 2)_= 38.36, *p* < 0.001), whereas no significant difference was observed between control and moderate drought treatments (Supplementary Fig. S3).

### Effects of drought stress and beet leaf miner infestation on leaf photosynthetic efficiencies

Drought stress and leaf miner infestation significantly affected the photosynthetic efficiencies in sugar beet leaves over time. The presence of drought stress resulted in a notable decrease in the maximum quantum yield of PS II (Fv/Fm) (LMM - drought: F_2,408_ = 601.66, *p* < 0.001). This reduction was particularly pronounced in plants exposed to high drought levels over an extended period (LMM - drought x time: F_6,408_ = 126.67, *p* < 0.001). Leaf mining also decreased the photosynthetic efficiency (LMM – beet miner: F_1,408_ = 7656.06, *p* < 0.001). Interestingly, this decrease was lowest in highly-stressed plants and most pronounced in moderately drought-stressed plants (LMM – beet miner x drought: F_2,408_ = 898.51, *p* < 0.001, Fig. 1A). Leaf miner infestation and drought also significantly reduced the effective photochemical quantum yield (Ф*_PSII_*) of PS II (LMM – beet miner: F_1,408_ = 2015.43, *p* < 0.001, drought: F_2,408_ = 252.94, *p* < 0.001, Fig. 1B) and again this effect was most pronounced at moderate drought conditions (LMM – beet miner x drought: F_2,408_ = 202.98, *p* < 0.001). A similar trend was observed for electron transport rate (ETR) (Supplementary Fig. S4). Detailed statistical outputs are provided in Supplementary Table S2.

**Fig. 1.**
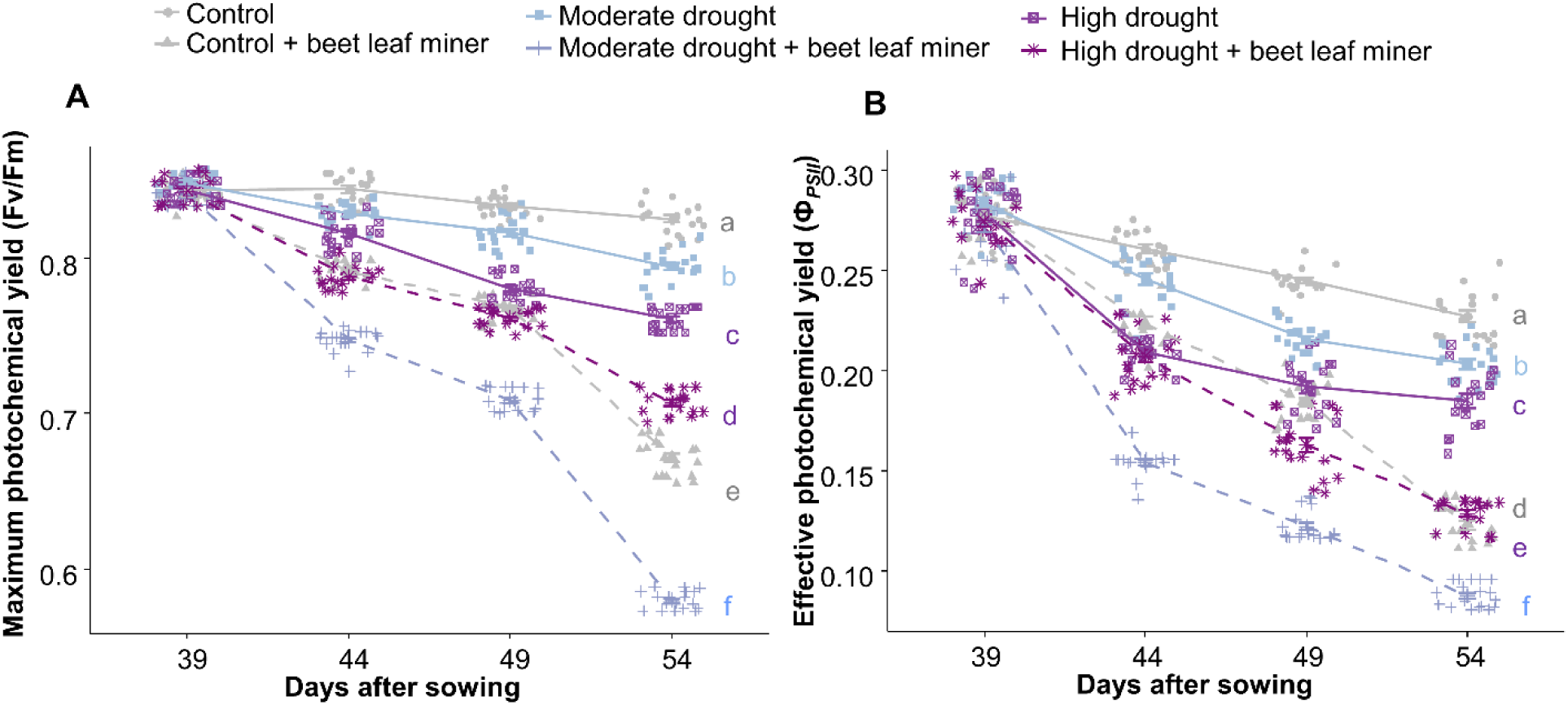
Effects of drought stress and leaf miner infestation on photosynthetic efficiency parameters in sugar beet leaves. (A) Maximum photochemical quantum yield of photosystem (PS) II (Fv/Fm) (LMM, n = 18). (B) Effective photochemical quantum yield of PS II (ФPSII) (LMM, n = 18). Data were analysed using linear mixed-effects models with drought (D; three levels: control, moderate, high), herbivory (H; present/absent), time (T), and their interactions as fixed factors. Significant main effects of D, H, and T, as well as significant interaction effects, were detected in all cases, as detailed in the supplementary section. Each data point represents an individual replicate. Different letters indicate statistically significant differences among treatments (p ≤ 0.05, n = 18).

### Effects of drought stress and beet leaf miner infestation on leaf central metabolites

A total of 114 metabolites were identified in sugar beet leaf tissue and classified into major metabolite groups, including amino acids, organic acids, fatty acids, and sugars (Fig. 2B) and visualized in a circular heatmap (Fig. 2A). Hierarchical clustering revealed five major metabolite groups based on their relative abundance (Fig. 2A), with concentrations generally increasing under drought and herbivory stress. However, some metabolites were significantly upregulated, and some were significantly downregulated based on the comparison of fold change (Supplementary Fig. S6 A-F) among different treatments.

**Fig. 2.**
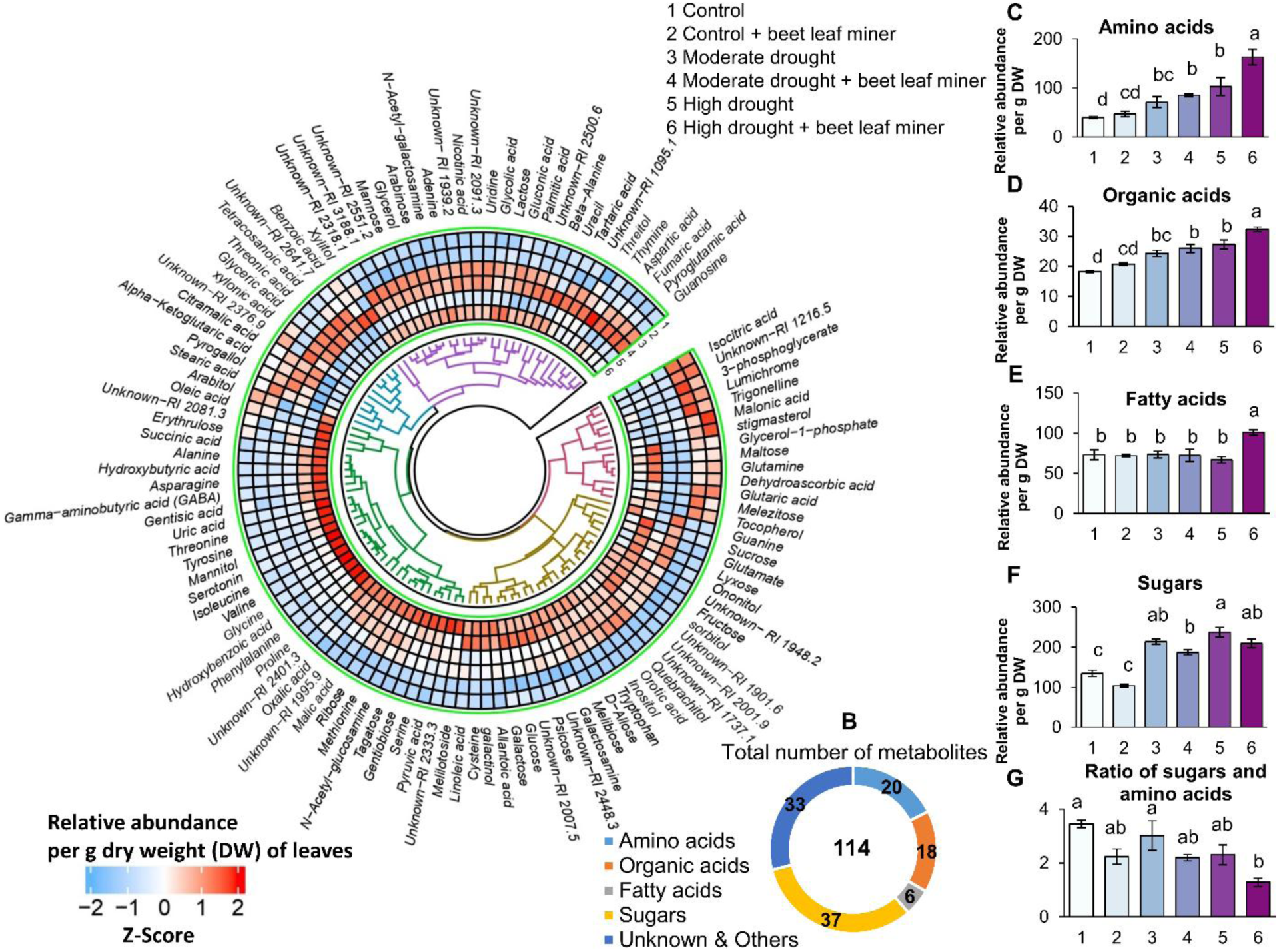
Effects of drought stress and leaf miner infestation on central metabolites in sugar beet leaves. (A) Circular heatmap with hierarchical clustering of all metabolites across different treatments. (B) Number of metabolites and derivatives categorized by major metabolic classes. Bar graphs represent approximate concentrations of (C) total amino acids (log10-transformed), (D) total organic acids, (E) total fatty acids, (F) total sugars, and (G) total sugar-to-amino acid ratio. Data were analysed using two-way ANOVA with drought (D; three levels: control, moderate, high), herbivory (H; present/absent), and their interaction (D × H) as fixed factors. Total amino acids were affected by D (p < 0.001) and H (p = 0.003), with no interaction (p = 0.27). Total organic acids showed main effects of D (p < 0.001) and H (p < 0.001) without interaction (p = 0.21). Total fatty acids were influenced by D (p = 0.045) and H (p = 0.01), with a significant D × H interaction (p = 0.001). Total sugars were affected by D (p < 0.001) and H (p < 0.001) without interaction (p = 0.97). The sugar-to-amino acid ratio decreased with D (p = 0.01) and H (p < 0.001), with no interaction (p = 0.94). Different letters indicate significance among treatments (p ≤ 0.05, n = 6). Data are presented as relative abundance, calculated as the peak area ratio of each analyte to the internal standard (adonitol).

Amino acid levels were significantly elevated in drought-stressed plants and by leaf miner infestation (LM: drought: F_2,30_ = 44.24, *p* < 0.001; leaf miner: F_1,30_ = 10.08, *p* = 0.003) (Fig. 2C). Organic acids followed a similar pattern with concentrations significantly affected by drought and leaf miner infestation (LM: drought: F_2,30_ = 52.06, *p* < 0.001; leaf miner: F_1,30_ = 14.14, *p* < 0.001) (Fig. 2D). In contrast, fatty acid levels were only increased in plants exposed to high drought combined with leaf miner infestation (LM: drought: F_2,30_ = 3.43, *p* = 0.045; leaf miner: F_1,30_ = 7.05, *p* = 0.01; drought × leaf miner: F_2,30_ = 8.67, *p* = 0.001) (Fig. 2E). Sugar metabolite concentrations were elevated under drought conditions but reduced in leaf miner treatments (LM: drought: F_2,30_ = 85.78, *p* < 0.001; leaf miner: F_1,30_ = 17.42, *p* < 0.001), with no significant difference between moderate and high drought levels (Fig. 2F). Lastly, the sugar-to-amino acid ratio declined with increasing drought stress and leaf miner infestation (LM: drought: F_2,30_ = 5.27, *p* = 0.01; leaf miner: F_1,30_ = 21.30, *p* < 0.001) (Fig. 2G).

Partial least squares discriminant analysis (PLS-DA) was conducted to identify stress-responsive metabolites in sugar beet. Metabolites with a variable importance in projection (VIP) score > 1 were considered significant, yielding 40 top-ranked compounds out of 120 total (Fig. 3A). To visualize overall patterns and treatment separation, dimensionality reduction was performed using principal component analysis (PCA) (Supplementary Fig. S7) and PLS-DA (Fig. 4B). In the PCA, PC1 and PC2 accounted for 39.1% and 19.8% of the variance, respectively, showing clear separation among treatments. The PLS-DA plot revealed clear clustering of treatment groups, indicating distinct metabolic profiles, suggesting that drought stress and beet leaf miner infestation led to significant metabolic changes.

**Fig. 3.**
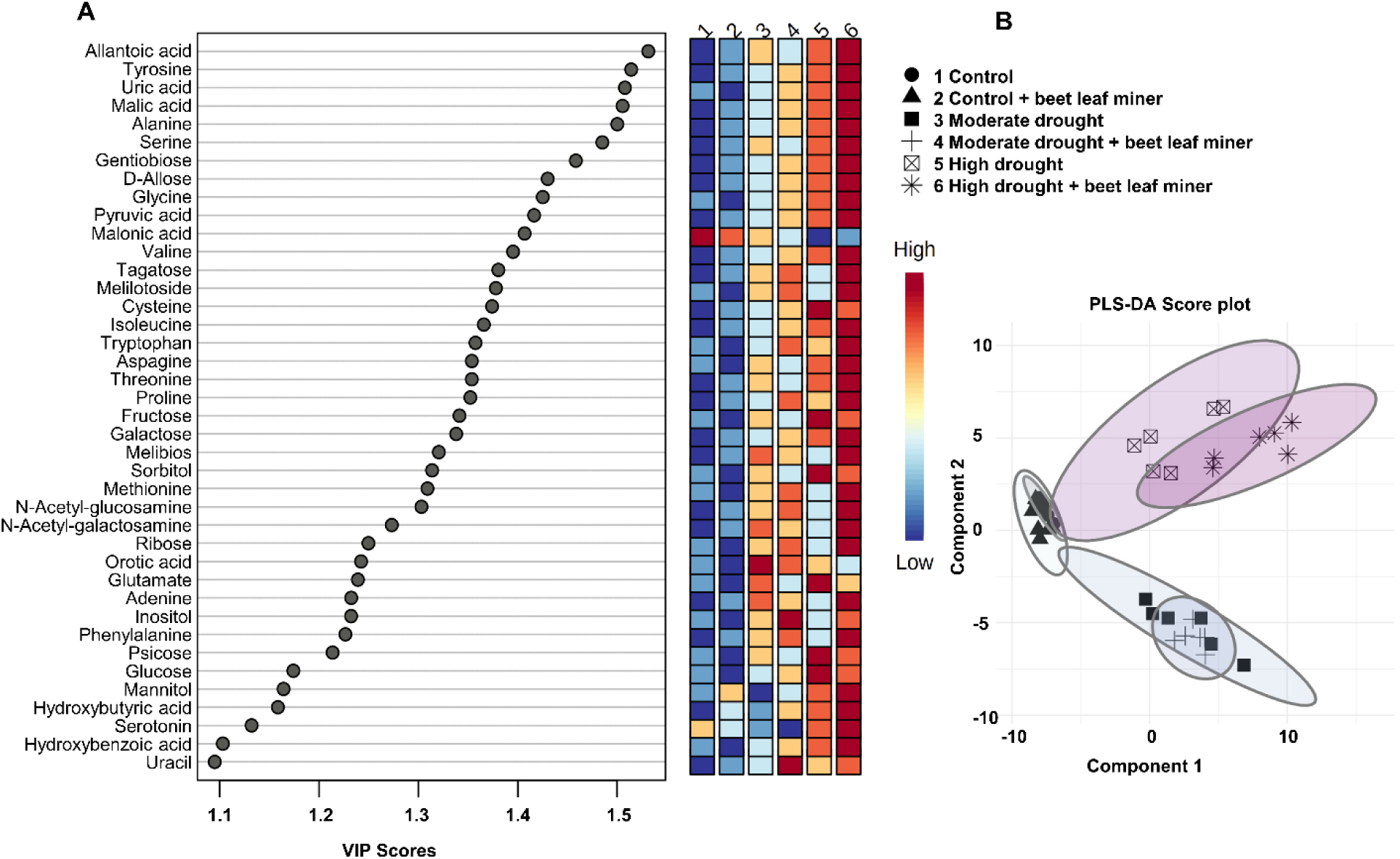
Important central metabolites in response to drought stress and leaf miner infestation were identified by partial least squares discriminant analysis (PLS-DA). (A) Forty top metabolites according to the variable importance of projection (VIP) score to different treatments, are shown. Colored boxes indicate the relative approximate concentrations of the corresponding metabolites in each group. (B) PLS-DA score plot based on their metabolite profiles.

**Fig. 4.**
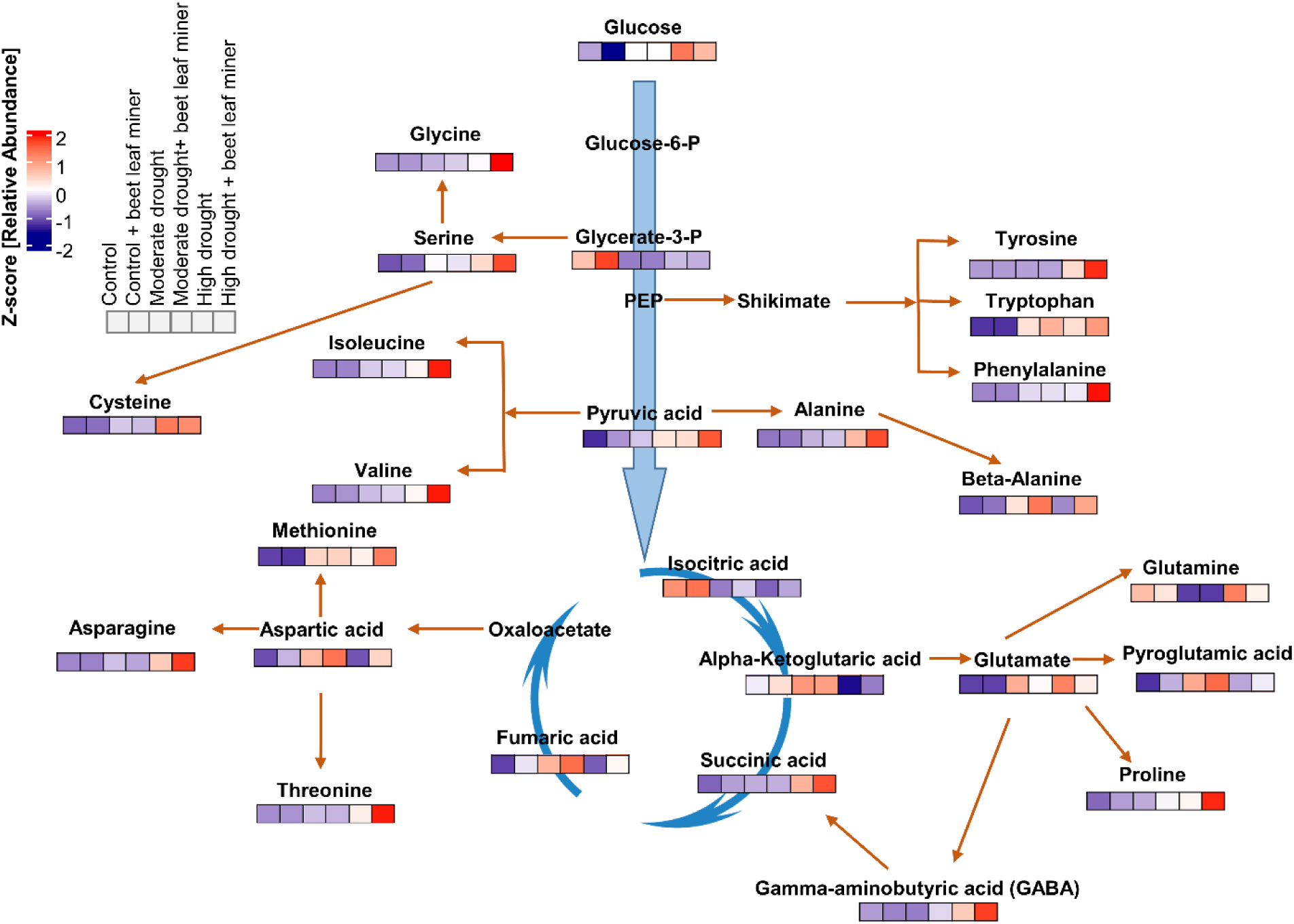
Metabolic changes involved in the amino acid pathway and tricarboxylic acid cycle under drought stress and beet leaf miner infestation. Each heatmap represents the normalized intensity of the corresponding metabolite; n= 6.

Based on VIP scores, the most responsive metabolites were primarily amino acids, along with selected organic acids and sugars. The concentrations of these metabolites generally increased with the severity of stress. Pathway analysis of amino acid biosynthesis revealed elevated levels of several key metabolites (Fig. 4), including a notable increase in glucose compared to control plants. Amino acids such as glycine, leucine, isoleucine, valine, proline, alanine, asparagine, threonine, and tyrosine showed significant increases relative to controls. Similarly, within the tricarboxylic acid (TCA) cycle, the intermediate succinic acid was more abundant in dual-stressed plants compared to plants exposed to one stressor (Fig. 4).

### Analysis of plant volatiles

Hierarchical clustering of qualitative and quantitative differences in VOC profiles revealed that beet leaf miner-infested control plants emitted a distinct blend compared to other treatments (Fig. 5A, Supplementary Table S3). When standardized by fresh weight (g FW), high drought + beet leaf miner plants released a more concentrated VOC blend than other treatments (Fig. 5B, Supplementary Table S4). Control plants emitted mostly green leaf volatiles (trans-2-hexenal, 3-hexenol, 1-octen-3-ol, octanal, cis-3-hexenyl-1-acetate) and monoterpenes (α-pinene, β-myrcene, 3-carene, D-limonene, β-ocimene, trans-β-ocimene, p-cymene, thymol, p-cymen-7-ol) at lower concentrations. Emission rate per plant tended to increase upon leaf miner infestation, but these differences were not significant. Some of these compounds (1-octen-3-ol, β-myrcene, trans-β-ocimene) were not emitted by drought-stressed plants, while the emission per plant of the other green leaf volatiles besides octanal was not affected by drought stress, but tended to increase upon herbivory, similar as in the controls. Most other compounds were absent from the VOC blends emitted by drought-stressed without herbivores. Sesquiterpenes—namely γ-elemene, β-guaiene, and carotol—were only detected in beet leaf miner-infested control and moderately drought-stressed plants. Notably, no sesquiterpenes were detected in high drought + beet leaf miner plants (Fig. 5A, B).

**Fig. 5.**
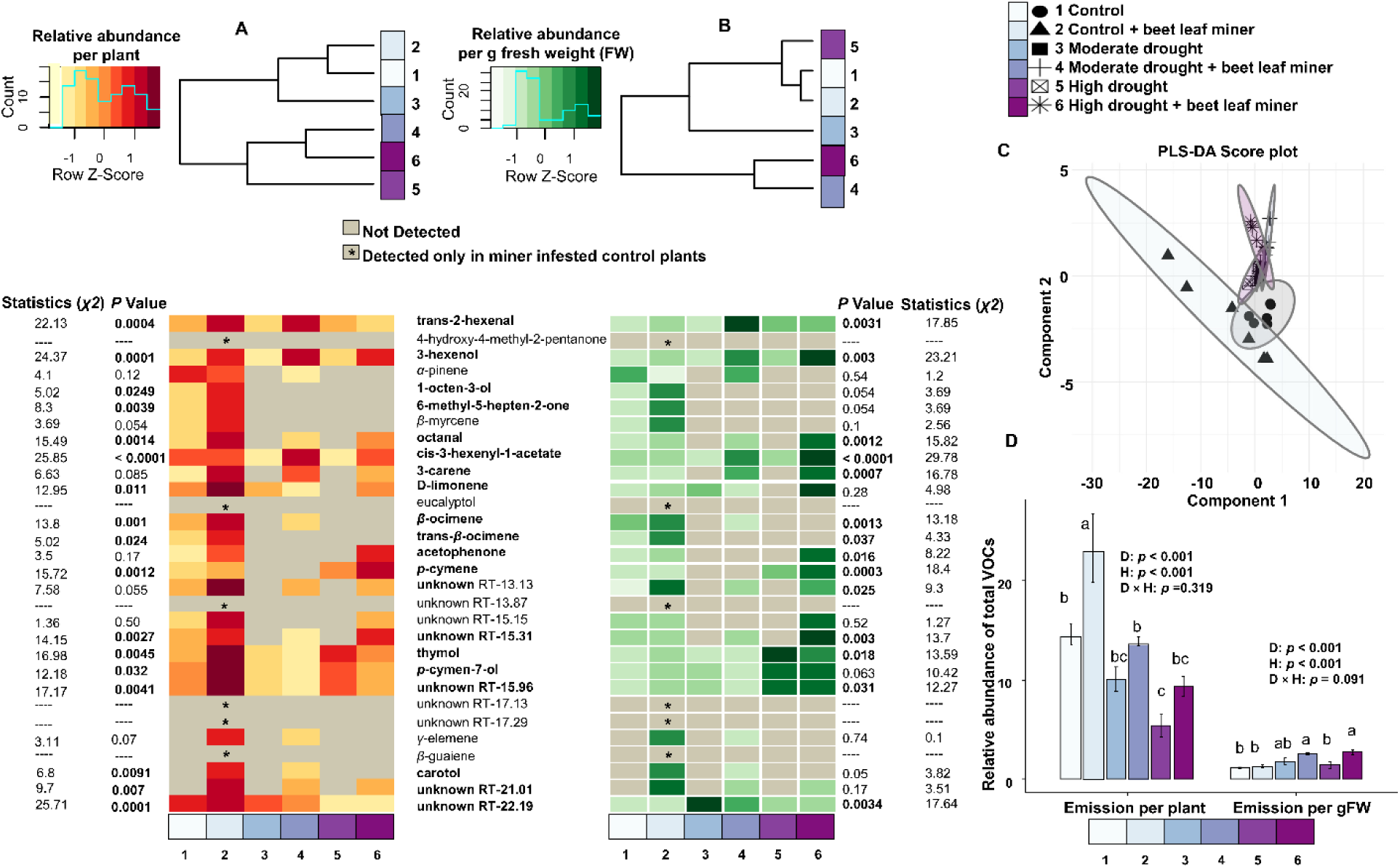
Heatmaps illustrate the emission rates of different VOCs from sugar beet plants. Kruskal-Wallis tests were used to compare emission rates of individual compounds between treatments. Significant differences between treatments for each compound are indicated alongside the heatmaps with χ² and P-values. Data are presented as relative abundances, calculated as the peak area ratio of each analyte to the internal standard (tetralin). Compound names and P-values in bold denote statistically significant differences among treatments (p ≤ 0.05, n = 6). (A) Sugar beet above-ground plant volatile emission from the whole plant. (B) Sugar beet above-ground plant volatile emission normalized per gram of fresh leaf weight. (C). PLS-DA score plot. (D) Total VOC emissions expressed both per plant and per gram of fresh leaf weight. Total VOC emissions (per plant and per gram fresh weight) were analysed using two-way ANOVA with drought (D; three levels: control, moderate, high), herbivory (H; present/absent), and their interaction (D × H) as fixed factors. Significance of main and interaction effects is indicated within panels.

To visualize overall patterns and treatment separation, dimensionality reduction was performed using principal component analysis (PCA) (Supplementary Fig. S8A) and PLS-DA (Fig. 5C). In the PCA, PC1 and PC2 accounted for 56.3% and 9.9% of the variance, respectively. The PLS-DA plot revealed clear clustering of treatment groups, indicating distinct VOCs profiles, suggesting that drought stress and beet leaf miner infestation led to significant VOCs changes. The top fifteen discriminatory compounds based on VIP scores are shown in Supplementary Fig. S8B.

Total VOC emission per plant increased upon herbivory (ANOVA: F₁,₃₀ = 15.67, p < 0.001) and decreased with increasing drought stress (ANOVA: F₂,₃₀ = 26.16, p < 0.001). When emission rate was standardized by fresh weight, opposite effects of drought stress were observed, because stressed plants were considerably smaller. VOC emission per g fresh weight increased with increasing drought stress (ANOVA: F₂,₃₀ = 9.02, p < 0.001) and herbivory in the drought-stressed plants (ANOVA: F₁,₃₀ = 12.39, p = 0.0013), while no increase in VOC emission upon herbivory was observed in the control (ANOVA: F₂,₃₀ = 2.59, p = 0.091) (Fig. 5D). Although high drought + beet leaf miner plants had the highest VOC concentration per gram FW, total emission per plant was significantly greater in control + beet leaf miner plants (Fig. 5D).

### Olfactory and oviposition responses of adult Pegomya cunicularia

In Y-tube olfactometer assays, beet leaf miner flies demonstrated a strong preference for VOCs emitted by sugar beet plants over soil substrate (Binomial GLM: *χ^2^* _(*df* 1)_ = 64.47, *p* < 0.001) (Fig. 6A). A significantly higher attraction was observed toward control plants compared to both high drought plants (Binomial GLM: χ²(1) = 92.92, p = 0.009) and high drought plants infested with beet leaf miners (Binomial GLM: χ²(1) = 86.85, p < 0.001). No significant differences were found among other odor pairs. In a choice test with different drought levels, female flies laid significantly more eggs on control plants (64.0 ± 1.6 eggs) than on moderate (15.8 ± 0.7 eggs) and high drought (4.4 ± 0.3 eggs) plants (GLMM: χ²(2) = 44.43, p < 0.001; control vs. moderate: p < 0.001; control vs. high: p < 0.001; moderate vs. high: p < 0.001) (Fig. 6B). Similarly, in a choice test including different levels of drought with and without beet leaf miners, oviposition was highest on control plants (40.8 ± 1.0 eggs), intermediate under moderate drought (20.0 ± 1.0 eggs), and lowest under high drought (7.9 ± 0.5 eggs, GLMM: drought χ²(4) = 226.43, p < 0.001; Fig. 6C). Plants infested with beet leaf miners received significantly fewer eggs across all treatments: control + beet leaf miner (13.5 ± 0.6), moderate drought + beet leaf miner (6.5 ± 0.7), and high drought + beet leaf miner (2.9 ± 0.4 eggs, GLMM: leaf miner χ²(1) = 53.5, p < 0.001; Fig. 6C). Egg numbers in a no-choice test were significantly affected by drought (GLMM: χ²(4) = 51.94, p < 0.001), and beet leaf miner infestation (GLMM: χ²(1) = 12.64, p < 0.001) (Fig. 6D). Under well-watered conditions, plants without beet leaf miner infestation had an average of 71 ± 2.1 eggs, whereas infested plants had 40.2 ± 1.8 eggs. High drought significantly reduced egg numbers to 35.2 ± 1.9 in uninfested plants and 15.3 ± 1.5 in infested plants (Fig. 6D), while no strong effect of drought stress was observed under moderate drought conditions.

**Fig. 6.**
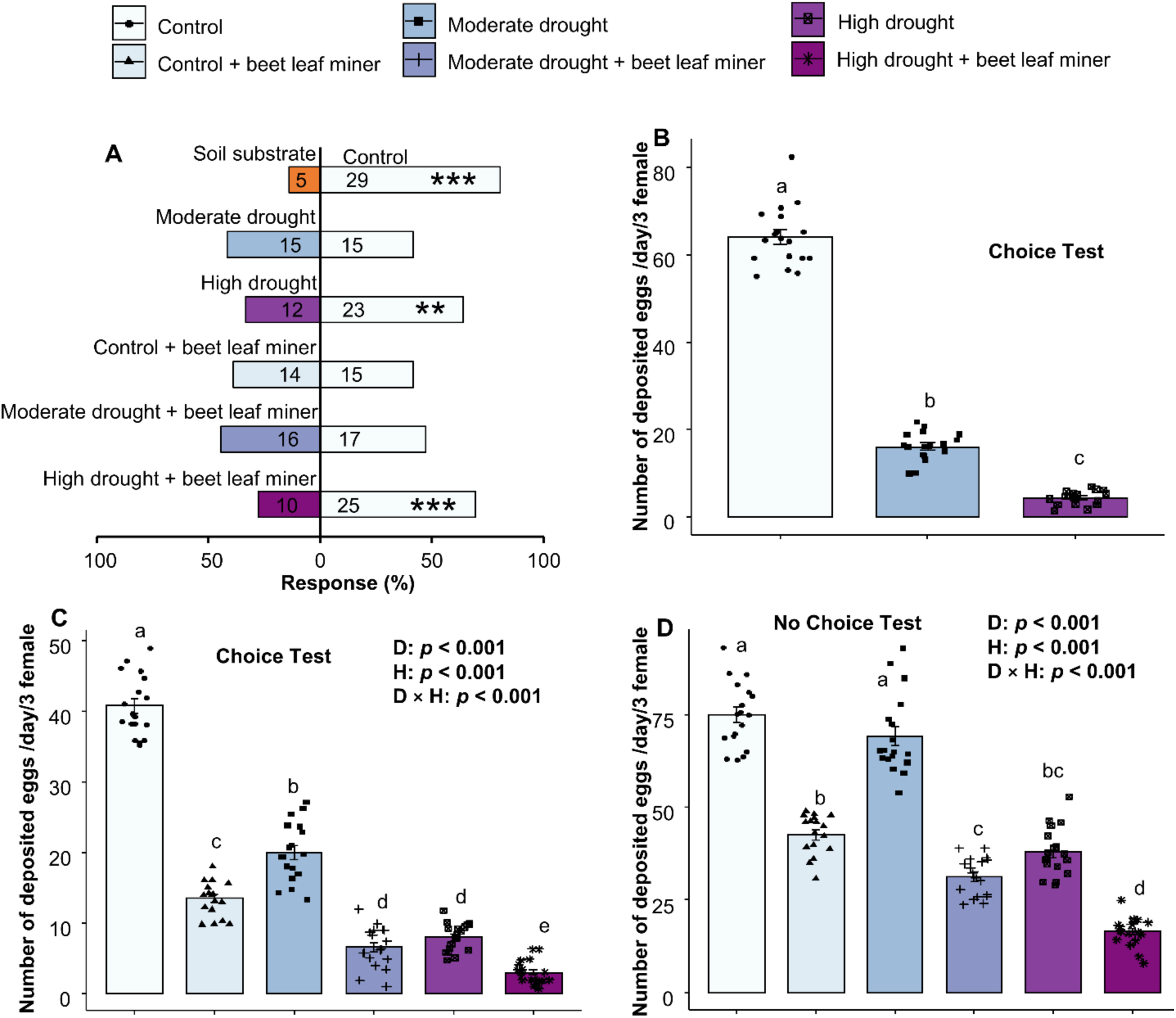
Olfactory and oviposition responses of beet leaf miner flies to sugar beet plants subjected to drought and leaf miner infestation treatments. (A) Y-tube olfactometer assay measuring the number of flies responding to two different odor sources at same time. Numbers inside the bars indicate the actual counts of responding flies (out of 36). Significance among treatments are indicated by asterisks (* p ≤ 0.05, ** p ≤ 0.01, *** p ≤ 0.001; Binomial GLM, n = 36). (B) Choice oviposition test with drought-only treatments presented simultaneously in a single tent (GLMM, p ≤ 0.05, n = 18). (C) Choice oviposition test including all treatments (drought and herbivore infestation) simultaneously presented in a single tent (GLMM, p ≤ 0.05, n = 18). (D) No-choice oviposition test with all treatments presented separately in individual tents (GLMM, p ≤ 0.05, n = 18). Oviposition data were analysed using generalized linear mixed models (GLMMs) with a Conway–Maxwell–Poisson (compois) distribution. Drought (D; three levels: control, moderate, high) and herbivory (H; present/absent) were included as fixed factors in both choice and no-choice tests. Significance of main and interaction effects is indicated within panels. Data points represent replicates. Different letters indicate statistically significant differences among treatments (p ≤ 0.05).

### Effects of drought stress on beet leaf miner development

Drought stress significantly affected the developmental success of *P. cunicularia*. The highest larval hatching rate from eggs was observed on moderately drought-stressed plants, whereas the lowest was recorded under high drought conditions (Binomial GLM: *χ^2^*_(*df* 2)_= 20.05, *p* < 0.001) (Fig. 7A). Correspondingly, feeding damage, quantified as mined leaf area (cm²), was greatest under moderate drought (Supplementary Fig. S5). When expressed as the proportion of mined leaf area, feeding damage was significantly higher in moderately drought-stressed plants compared to both control and severe drought treatments (Kruskal-Wallis: *χ^2^*_(*df* 2)_= 41.54, *p* < 0.001) (Fig. 7B). Moderate drought conditions also favored *P. cunicularia* development by leading to highest pupal (LMM: F_2,14_= 9.30, *p* = 0.002) and adult (LMM: F_2,14_= 13.53, *p* < 0.001) weights (Fig. 7C). Furthermore, developmental timing was affected: insects took significantly longer to develop under moderate drought conditions. This included egg-to-pupa duration (Kruskal-Wallis: *χ^2^*_(*df* 2)_= 61.98, *p* < 0.001), pupa-to-adult emergence time (LMM: F_2,14_= 16.10, *p* < 0.001), and total development time (Kruskal-Wallis: *χ^2^*_(*df* 2)_= 34.38, *p* < 0.001) (Fig. 7D).

**Fig. 7.**
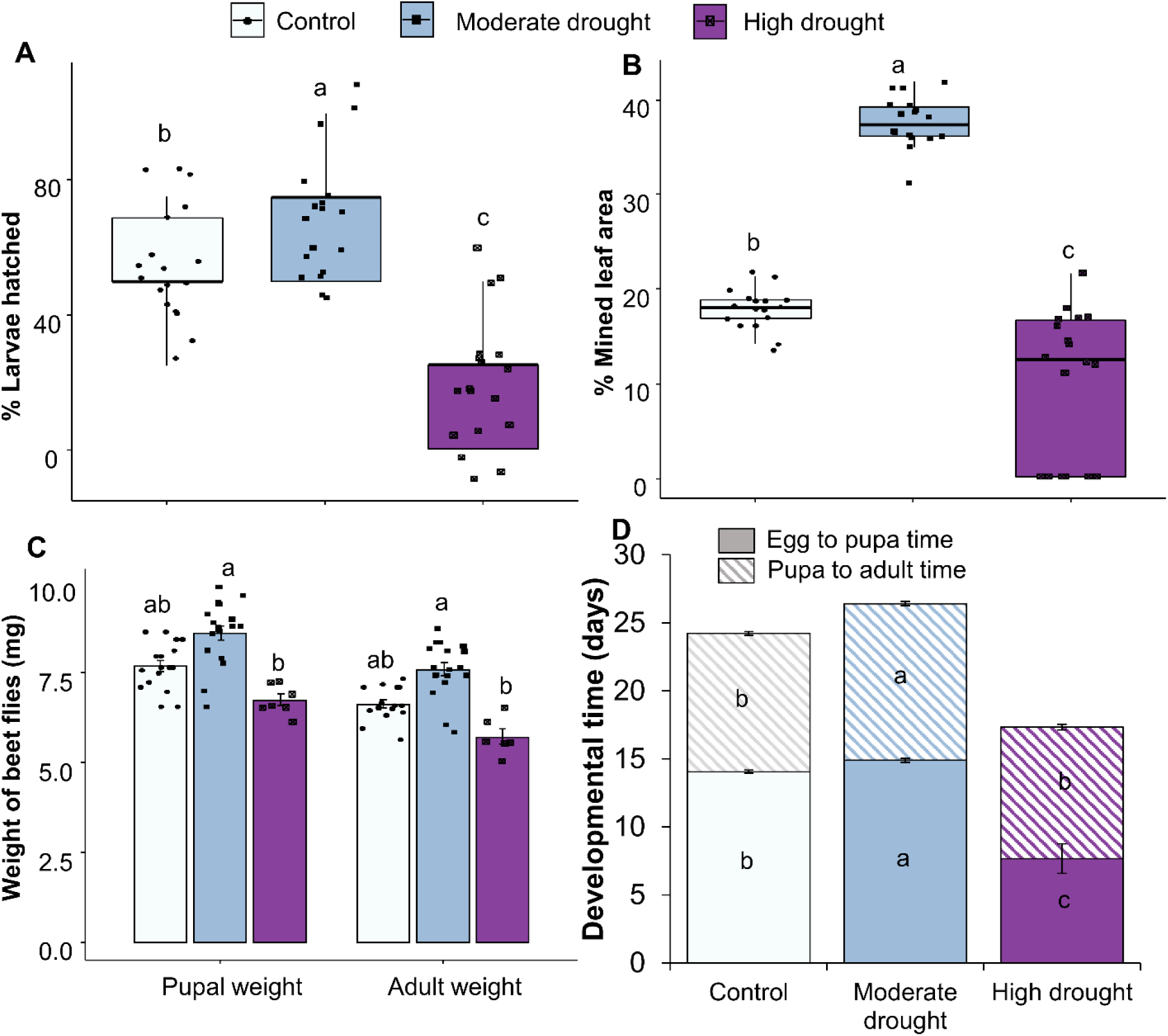
Effects of varying drought levels on the performance of *Pegomya cunicularia*. (A) Percentage of hatched larvae (Binomial GLM, n = 18) (A). (B) Percentage of mined leaf area (Kruskal–Wallis, n = 18). (C) Pupal and adult weight (LMM, n = 18). (D) Developmental time including egg-to-pupa and pupa-to-adult durations (Kruskal–Wallis, n = 18). Each data point represents an individual replicate. Different letters indicate statistically significant differences among treatments (p ≤ 0.05).

## Discussion

Climate change leads to more frequent drought events and increased herbivore pressure on crop plants, often occurring simultaneously. Our results demonstrate that the combined effects of drought and herbivory by the beet leaf miner, *P. cunicularia*, elicit profound, non-additive physiological and metabolic reprogramming in sugar beet, exceeding the impact of either stressor alone. Notably, drought coupled with leaf miner infestation triggered a complex pattern: enhanced nutritional quality of leaf tissues supported superior larval performance under moderate drought conditions, while the concurrent suppression of VOC emissions reduced plant apparency and decreased oviposition by adult flies, particularly under high drought.

Consistent with established drought acclimation strategies (Yang et al., 2021; Kurepa and Smalle, 2022), sugar beet exposed to drought exhibited reduced plant height, shoot biomass, and leaf area, alongside a higher root-to-shoot ratio (Figure S2 A-J), likely reflecting adaptive prioritization of root growth to improve water uptake. Such structural changes are known to be tied to stomatal closure and hormonal signaling cascades modulated by reactive oxygen species (ROS) and calcium fluxes (Zebelo and Maffei, 2015; Agurla et al., 2018). Importantly, moderate drought did not significantly diminish leaf water content (Figure S3), thus maintaining turgor sufficient to support herbivore feeding, whereas severe drought lowered tissue hydration and biomass more drastically, indicating a physiological threshold beyond which plant performance and palatability decline.

Photosynthetic parameters (Fv/Fm, ΦPSII) were significantly impaired by both drought and herbivory, with the most pronounced and synergistic reductions observed under combined moderate drought and beet leaf miner infestation (Figure 1). This pattern likely reflects enhanced herbivore performance on moderately drought-stressed plants, where greater leaf miner feeding damage was observed. The resulting intensified tissue injury, presumably coupled with elevated ROS accumulation and exacerbated chloroplast structural damage and photoinhibition, is consistent with reports from drought-stressed cotton and herbivore-challenged maize (Deeba et al., 2012; Shin et al., 2021; de Souza et al., 2020). Conversely, under severe drought, the constrained physiological activity of both plants and herbivores may have limited further damage, leading to less pronounced additional photosynthetic decline despite water stress severity. Our results suggest that moderate drought sensitizes photosynthetic machinery to insect-induced damage even when water limitation is not critical, supporting the idea that abiotic stress may exacerbate biotic damage via oxidative stress pathways.

As outlined above, *P. cunicularia* developed best under moderate drought intensity, with beet miner larvae exhibiting the highest larval hatching rates, pupal and adult weights, and prolonged developmental time (Figure 7). This enhanced performance resulted in greater feeding damage on moderately drought stressed plants. These findings align with reports that moderate drought can improve tissue nutritional quality for leaf miners (Hahn and Maron, 2018; Santiago-Salazar *et al*., 2022). In contrast, high drought suppressed larval development and survival, indicating a physiological threshold beyond which stress impairs herbivore performance. Prior studies showed that drought can increase nutrient content in host tissues, thereby enhancing herbivore performance—but only within a specific intensity window (Franzke and Reinhold, 2011; Sconiers and Eubanks, 2017; Sconiers *et al*., 2020). Our data support this model, implying that the moderate drought condition created a nutrient-rich, developmentally favorable environment for the beet leaf miner.

Metabolic profiling of sugar beet leaves revealed alterations in central carbon metabolism, amino acid levels, and specialized metabolism in response to combined drought and beet leaf miner stress. For instance, elevated levels of amino acids such as tyrosine, alanine, and serine (highlighted by their high VIP scores) may reflects shifts in stress-adaptive pathways, including nitrogen recycling and antioxidant defense (Rocha et al., 2010; Maeda and Dudareva, 2012; Igamberdiev and Kleczkowski, 2018). The observed upregulation of aromatic amino acids and derivatives (e.g., dopamine) suggests modulation of specialized metabolism involved in plant defense (Kulma & Szopa, 2007). Meanwhile, fluctuations in TCA cycle intermediates reflect dynamic energy and nitrogen rebalancing under multi-stress conditions (Obata & Fernie, 2012). These combined metabolic shifts underscore the plant’s integrated and flexible biochemical response to complex environmental challenges, with synergistic stress treatments producing the most pronounced effects (Krasensky & Jonak, 2012; Atkinson & Urwin, 2012).

Beyond nutrient provisioning, amino acid levels may also influence central carbon metabolism and defense-related secondary pathways (Figure 4). Changes in amino acid and TCA intermediates under combined stress could contribute to volatile biosynthesis, although these changes may also reflect altered conjugation, turnover, or compartmentalization rather than de novo biosynthesis. The extensive reprogramming of amino acid and organic acid metabolism in response to drought and insect herbivory might therefore affect VOC production, as many volatiles are derived from central metabolic precursors (Pott *et al*., 2019). Female herbivorous insects rely on these volatile cues to identify and select suitable host plants for oviposition (Achhami *et al*., 2021; De Moraes *et al*., 2001).

When sugar beet plants were attacked by leaf miners, they responded, like many other plants under herbivore pressure, by increasing the emission of plant volatiles (Guerrieri and Rasmann 2024). This response is triggered by the plant’s detection of herbivore damage, involving both the specific feeding patterns of the insect and the presence of chemical elicitors in the insect’s saliva. Recognition of these cues activate the jasmonic acid defense pathway, which in turn stimulates the expression of genes for VOC biosynthesis (Erb and Reymond, 2019; Heil, 2014). In contrast, reduced water availability was associated with lower VOC emissions per individual plant. It might appear intuitive to assume that drought-induced stomatal closure was partly responsible for the reduced VOC emissions (Lin *et al*., 2022). Drought conditions limit stomatal opening, which constrains the instantaneous release of VOCs (Loreto and Schnitzler, 2010; Brilli *et a*l., 2007). However, in our study, drought also caused a reduction in total plant biomass, resulting in smaller VOC emitting plants. When emissions were normalized per unit of plant biomass, VOC production was not reduced. In fact, sugar beets experiencing both drought and herbivore attack released even higher VOC amounts per unit biomass than well-watered counterparts. This indicates that the biochemical capacity for VOC biosynthesis within the plant tissue remained unaffected or was upregulated despite physiological constraints on volatile release from stomata. Importantly, emission rates expressed per unit biomass reflect metabolic regulation at the tissue level, whereas herbivores in natural settings respond to whole-plant odour plumes. Thus, total VOC emission per plant represents the ecologically relevant signal available for host location. Under drought conditions, reductions in plant size can therefore decrease the overall odour signal even if biosynthetic capacity per unit tissue is unchanged or enhanced. Consequently, interpretation of insect behavioural responses should primarily consider whole-plant emission rates, while biomass-normalized values provide insight into underlying physiological adjustments. A similar pattern was also observed by Rahman *et al*. 2025: total VOC emission per plant from aphid-infested sugar beet was strongly reduced upon severe drought stress, while VOC emission per unit biomass was not. Our findings align with those observed in maize subjected to herbivory and salinity stress, which shares similarities with drought stress, where VOC emissions were reduced per plant but remained unchanged when normalized to biomass (Forieri *et al*., 2016). Furthermore, these results also align with the higher concentrations in central metabolites subjected to the combined stresses.

Apart from differences in total emission rates, sugar beet plants also showed unique VOC profiles (Figure 5). VIP score analysis pinpointed several key compounds, such as cis-*β*-ocimene, *β*-myrcene, 1-octene-3-ol, trans-*β*-ocimene, *β*-guaiene, eucalyptol, and 4-hydroxy-4-methyl-2-pentanone, as principal contributors shaping the structure of these blends (Supplementary Figure S8). Plant volatiles serve as critical chemical cues that guide herbivores in locating suitable host plants for oviposition. They provide information about plant identity and stress status, enabling the insect to select optimal sites for egg laying. Changes in VOC profiles due to abiotic stresses or insect feeding can influence herbivore behavior, affecting host selection and ultimately plant-insect interactions (Bruce *et al*. 2005; Bruce and Pickett 2011). Our Y-tube experiments confirmed that adult beet flies preferred the odor of uninfested, well-watered and moderately drought-stressed plants, while they disliked the VOC blend of severely drought-stressed sugar beet. The drought-induced suppression of key volatiles may have reduced plant attractiveness to flies. Interestingly, the VOC blends induced by leaf mining from conspecific larvae neither increased nor decreased the attractiveness of sugar beets compared to undamaged plants, which contrasts with their oviposition behavior.

When *P. cunicularia* females were allowed to assess healthy and stressed sugar beet plants using multiple sensory cues, including but not limited to olfaction, in both choice and no-choice experimental setups, distinct oviposition preferences emerged. In comparative assays between well-watered and drought-stressed plants, females showed a strong preference for well-watered sugar beet. Additionally, the presence of conspecific larvae mining leaves significantly reduced egg deposition on these plants. Notably, drought and leaf mining exhibited strong negative synergistic effects on oviposition choice. However, in no-choice situations where females could not compare plants under different stress treatments, moderate drought alone did not discourage egg laying. This latter scenario likely mimics agricultural conditions where sugar beet monocultures experience uniform drought stress across entire fields.

The “mother knows best” hypothesis postulates that female insects choose oviposition sites that maximize the survival and fitness of their offspring (Jaenike 1978; Jones 2022). Our results support this hypothesis, as *P. cunicularia* was not attracted to the VOCs plants und severe drought stress and laid fewer eggs on them. Larvae developing on these highly stressed plants exhibited reduced feeding and the resulting adults were significantly smaller. Interestingly, the comparison between well-watered and moderately drought-stressed sugar beet was less straightforward. More larvae hatched and fed on moderately stressed plants, and although the difference was not statistically significant, they tended to be heavier at the adult stage. However, female flies never preferred moderately drought-stress plants, neither in the olfactometer nor for oviposition; at best these plants were equally attractive as well-watered plants.

In the oviposition assays, plants were not enclosed and females had unrestricted access to plant surfaces, allowing multiple sensory modalities to contribute to host selection. Therefore, in addition to volatile cues, visual signals, leaf surface properties, and contact chemical cues may have influenced oviposition decisions (Rojas et al., 2003; Fernández et al., 2019; Saitta et al., 2024). The Y-tube olfactometer assays, which isolate airborne cues, demonstrated that drought and herbivory altered volatile-mediated orientation behaviour. The oviposition assays further indicate that these volatile-driven preferences translate into realized host selection under more natural, multimodal conditions. Together, the combination of olfactometer and oviposition experiments strengthens the conclusion that drought-modulated plant responses influence host selection behaviour, with volatile cues likely acting at longer range and contact cues potentially refining final acceptance decisions.

Elevated levels of plant sugars and amino acids are known to create nutrient-rich conditions that favor herbivore growth and development (Li et al., 2024). Thus, based solely on the observed accumulation of central metabolites, one might predict optimal herbivore performance under high drought conditions, where nutrient concentrations were most elevated. However, despite high metabolite concentrations under high drought, herbivore performance was optimal under moderate drought, suggesting that enrichment of central metabolites alone does not determine host suitability. As leaf water content was comparable between control and moderate drought plants but significantly reduced under high drought (Supplementary Fig. S3), we infer that tissue hydration modulated the bioavailability and palatability of accumulated nutrients (Loomis, 1997; Wei et al., 2000). Furthermore, high drought may have increased the concentration of defensive metabolites, thereby offsetting the potential benefits of increased nutrient levels. While our analysis focused on central metabolites and volatile organic compounds, drought is also known to influence non-volatile specialised metabolites involved in plant defense, such as phenolics and other compounds. These were not explicitly quantified in the present study, as our primary aim was to link central metabolic reprogramming and volatile-mediated signaling to multi-trophic interactions. Nevertheless, stress-induced shifts in primary metabolism may affect downstream defensive pathways, potentially contributing to the observed patterns in herbivore performance and oviposition behaviour. Although high drought caused the greatest overall increase in central metabolite concentrations, the metabolite profiles of plants under moderate and high drought stress were distinctly different. Whether these differences in metabolite quality translate into variations in nutritional value for *P. cunicularia* remains to be tested in feeding experiments.

In conclusion, our study demonstrates that combined drought and herbivory induce complex physiological and metabolic changes in sugar beet, profoundly affecting both plant traits and herbivore dynamics. Moderate drought improves larval performance by enhancing leaf tissue nutritional quality, while simultaneously altering volatile organic compound (VOC) emission profiles that serve as important chemical cues for the leaf miner *P. cunicularia* in host plant selection. Rather than acting as a defense mechanism, these VOC changes convey information about plant stress status, influencing adult oviposition behavior and leading to reduced egg laying on highly drought-stressed plants. These findings highlight the nuanced interactions between abiotic stress, plant metabolic status, and herbivore host-finding strategies. Given the increasing frequency of drought events under climate change, understanding how environmental factors modulate plant chemistry and herbivore behavior is critical for developing pest management approaches that consider both plant physiology and insect ecology in agroecosystems.

## Acknowledgements

We thank Dr. Bernd Ulber for collecting the first sugar beet flies (*P. cunicularia*) from the field and Dr. Verner Michelsen (Associate Professor Emeritus, The Natural History Museum, Copenhagen) for species identification. We also thank Jonas Watterott for technical support and Dr. Franz Hadacek for optimizing the GC-MS method for central metabolite analysis and helpful comments on the manuscript.

## Author contributions

SR: Conceived and designed the study, performed experiments, collected and analyzed data, and drafted the manuscript.

MZS: Conducted experiments, performed metabolome data analysis, and reviewed the manuscript.

IV: Contributed to study conceptualization, critically reviewed and edited the manuscript, and supervised the research.

MR: Conceived the study, critically reviewed and edited the manuscript, and supervised the research.

## Data availability

All data supporting the findings of this work are available within the paper and its supplementary data.

## Supplementary data

### Methodology for artificial mass rearing

#### Plant, insect, and rearing condition

Six-week-old sugar beet plant (*Beta vulgaris*) subspec. *vulgaris* cultivar ‘Vasco’ (SESVanderHave, Belgium) was used as a host. Beet flies *Pegomya cunicularia* were collected in May 2020 from the sugarbeet field in Göttingen, Germany. The rearing room condition was 16L:8D photoperiod, light intensity 130 ± 10 µmol/(s m^2^), relative humidity of 65 ± 5 %, and a temperature of 20 ± 2 °C.

#### Artificial diet for the flies

Two types of food were supplied at the same time i) Dry food: prepared with skimmed milk powder, dextrose, soya meal, and yeast with the ratio of 10:10:1:1, and ii) Wet food: prepared with honey, soya meal, and yeast with the ratio of 5:5:1 and mixed with a drop of water to make it creamy. Water was also provided in a separate glass vial to the flies, and fresh foods were provided every seven days.

#### Rearing technique

Three separate tents (60 × 60 × 70 cm) were used for successful rearing. One tent was used for oviposition, containing sugar beet plants, beet flies (male and female), and food for the flies. Generally, a single female fly can lay 20-30 eggs/day, enough for a single plant to rear the larval stage. Adult flies were allowed to oviposit for 24 h, after which the plants were transferred to the second tent to allow larval development. Water was sprayed daily on the plants with developing larvae for better performance of the beet leaf miner. The larval period generally takes 10-15 days, and before finishing the larval period (10 days), leaves were cut at the base of the petiole and placed in a tray (10 × 20 × 8 cm), containing a mixture of soil substrate (sand:clay:organic matter; 2:1:1) and transferred to the third tent for pupation. It generally takes 10-15 days until adult emergence, and pupae were maintained by daily spraying water on the soil substrate. Finally, the adult flies were transferred to the first oviposition tend to continue the rearing process (Supplementary Fig. S1).

### Statistical analysis

#### Assessment of model assumptions

For all linear models (LM) and linear mixed-effects models (LMM), assumptions of normality of residuals and homogeneity of variance were evaluated prior to inference. Normality was assessed using visual inspection of Q–Q plots and residual histograms, and homoscedasticity was examined by plotting residuals against fitted values. For generalized linear (mixed) models (GLM/GLMM), model fit and distributional assumptions were evaluated using diagnostic simulations. When assumptions of normality and/or homoscedasticity were not met, data were either appropriately transformed (e.g. log₁₀ transformation) or analysed using generalized linear mixed models with suitable error distributions. In cases where model convergence was not achieved or model fit was inadequate due to data structure or limited sample size, non-parametric tests (Kruskal–Wallis followed by Dunn’s test) were applied.

##### Experiment 1

Three drought boxes with six plants each per treatment were used to monitor the morphological features of sugar beet plants, resulting in a total of n = 18 plants for each of the treatments (control, moderate drought, high drought). A generalized linear mixed effect model (GLMM) with a log link to Gaussian family distribution was used for analyzing changes in sugar beet plant height over time. Linear mixed models (LMMs) with drought and time as fixed factors and box as random factor were used to analyze total biomass, root weight, shoot weight, root shoot ratio by weight, root length, and average leaf area, by using the package “nlme” (José Pinheiro *et al*., 2023). Shoot length, number of leaves, and sugar beet leaf water content were analyzed by Kruskal-Wallis non-parametric test. Root shoot ratio by length was analyzed by generalized linear mixed effect model (GLMM) with “glmmTMB” (Brooks *et al*., 2017) function followed by Gaussian family distribution. Model assumptions were checked by “DHARMa” (Florian Hartig, 2022) package. Tukey HSD posthoc tests were used after GLMMs and LMMS by using “emmeans” (Lenth *et al*., 2023) and “multcomp” (Hothorn *et al*., 2008) packages. The Dunn test from the “rcompanion” (Mangiafico, 2023) package was used as posthoc test after the Kruskal-Wallis tests.

##### Experiment 2

Six drought boxes were used, one for each treatment (control, control + beet leaf miner, moderate drought, moderate drought + beet leaf miner, high drought, and high drought + beet leaf miner). Each box contained six plants, and three boxes were assigned per treatment (total n = 18 plants per treatment). Drought was applied at the box level, so each box represented one block (experimental unit), and plants within a box experienced the same conditions. Chlorophyll fluorescence parameters such as the maximum photochemical quantum yield of PSII (Fv/Fm), the effective photochemical quantum yield of PSII (ΦPSII), and the electron transport rate (ETR) were measured for each plant. Linear mixed models (LMMs) were used to analyze the data, with drought, beet leaf miner, and time as fixed factors, and plant identity nested within block as a random factor. The effect of drought stress on the number of larvae that hatched from the eggs was analyzed by GLMM with binomial family distribution, including block as a random effect. Continuous traits- including mined leaf area, pupal and adult weight, and pupae to adult time were analyzed by LMMs, with block as a random factor. Tukey HSD tests for multiple comparisons were used as posthoc tests. Percent mined leaf area, egg-to-pupal time, and total development time did not meet the assumptions of normality and homoscedasticity required for linear (mixed) models. We therefore considered generalized linear mixed models (GLMMs) with appropriate distributions (binomial/beta for percent data, log-normal for development times). However, these models either failed to converge or provided poor fits due to the limited sample size and data structure. Consequently, we applied the non-parametric Kruskal–Wallis test, which does not assume normality, followed by Dunn’s test for pairwise comparisons.

##### Experiment 3

Metabolomics data sets were obtained from leaf samples (n= 6). A linear model was employed (LM) to analyze the effect of drought on total amino acid, organic acid, fatty acid, and sugar metabolites and the sugar-to-amino acid ratio. Subsequently, a two-way ANOVA was conducted, and multiple comparisons were performed using the Tukey HSD posthoc test. The relative concentration of amino acid was log_10_-transformed before the statistical analysis. Circular heatmap and clustering representation of all metabolites were analyzed with software R with Z-score initialization. Both supervised partial least squares-discriminant analysis (PLS-DA) and unsupervised principal component analysis (PCA) were performed in R to reduce dimensionality and visualize relationships among treatments. For VIP score calculation, data were normalized by log10 transformation and auto-scaling (mean-centered and divided by the standard deviation of each variable). The top forty compounds were then identified based on VIP scores using MetaboAnalyst 5.0 (https://www.metaboanalyst.ca/).

##### Experiment 4

Hierarchical clustering was computed for VOCs based on the length of the straight line drawn followed by Euclidean distance with complete linkage by using software R. The influence of drought stress and herbivory on the emission of individual VOCs was analyzed with Kruskal-Wallis non-parametric test (n= 6). 2-way ANOVAs with drought and herbivory as factors were performed to analyze total VOC emission per plant, and per g FW of leaves, followed by Tukey HSD posthoc test for multiple comparisons. Principal component analysis (PCA) and partial least square-discriminate analysis (PLS-DA), of VOCs profiles of different treatments were analyzed using software R.

##### Experiment 5

In the Y-tube olfactometer experiments, the responses of *Pegomya* beet flies to the two odor sources were analyzed using a generalized linear model (GLM) with a binomial error distribution. The response variable was specified as a two-column vector (cbind(odor1, odor2)), representing the number of flies choosing each odor source. Flies that did not make a choice were excluded from the analysis. The total number of flies tested and the proportion of non-responders are reported in result section.

##### Experiment 6

The number of eggs laid by *Pegomya* flies were counted in the choice and no-choice test (n=18). The effect of drought-stress and the presence of conspecific larvae on oviposition preferences was analyzed by generalized linear mixed models (GLMM) with Conway-maxwell-Poisson (compois) family distribution from the “glmmTMB” package (Brooks *et al*., 2017). Drought and herbivory were included as fixed factors (for both choice and no-choice test), and tent was added as a random factor only in the no-choice test. Multiple comparisons were performed with Tukey HSD post hoc tests using the “emmeans” (Lenth et al., 2023) and “multcomp” (Hothorn et al., 2008).

---

**Supplementary Fig. S1:**
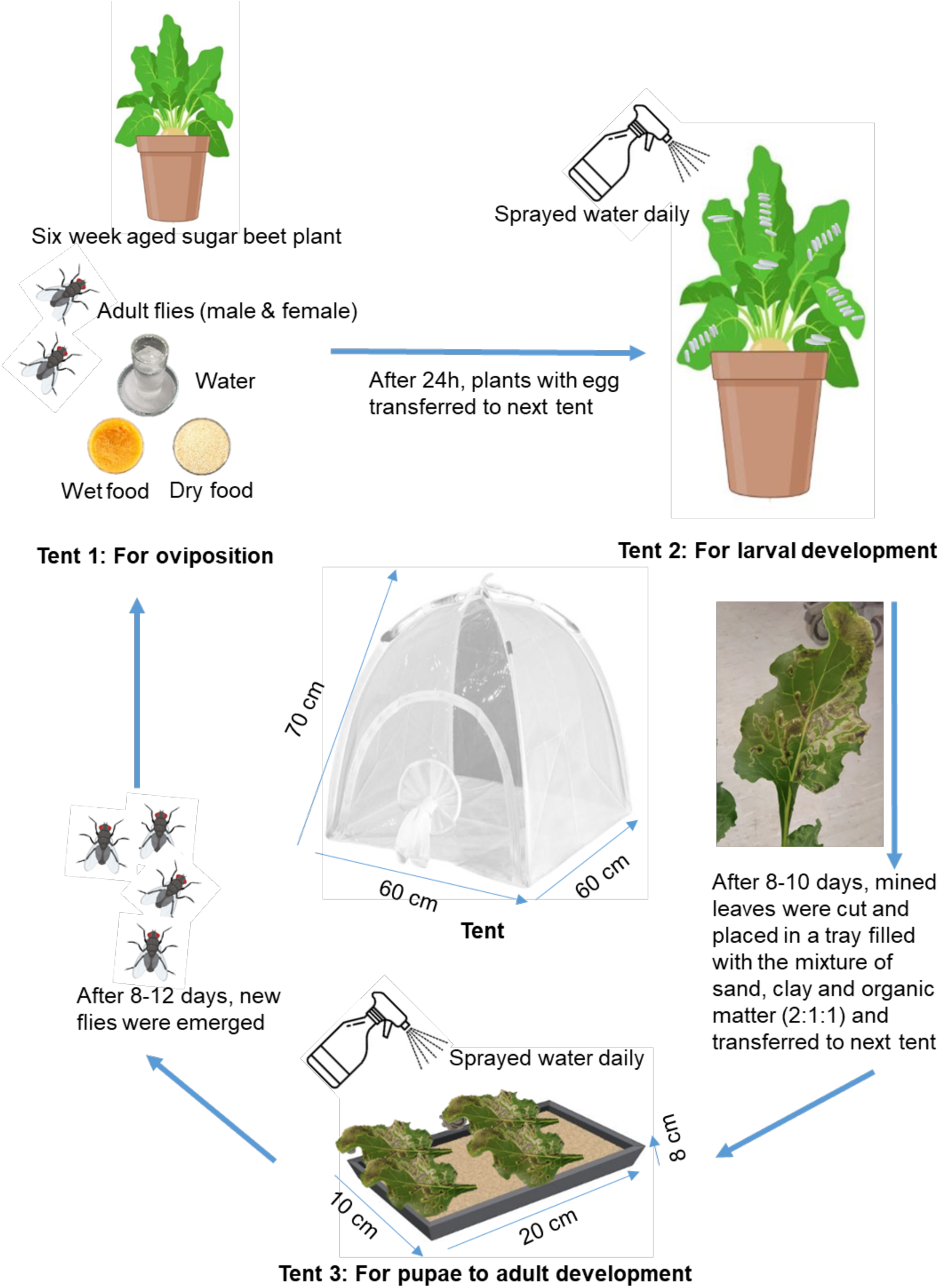
Schematic representation of laboratory method for the rearing of sugarbeet leaf miner *Pegomya cunicularia*.

**Supplementary Table S1:** Compound detected among different treatments summarized with their retention index from experiment and literature (NIST database)

| Compound name | Retention Index<br>Experimental | Retention Index<br>Literature |
| --- | --- | --- |
| trans-2-hexenal | 801.26 | 822 |
| 4-hydroxy-4-methyl-2-pentanone | 845.54 | 841.5 |
| 3-hexenol | 856.89 | 858 |
| α-pinene | 936.45 | 937 |
| 1-octen-3-ol | 980.61 | 983 |
| 6-methyl-5-hepten-2-one | 989.05 | 998 |
| β-myrcene | 991.88 | 992 |
| octanal | 1004.5 | 1004 |
| cis-3-hexenyl-1-acetate | 1007.65 | 1007 |
| 3-carene | 1013.31 | 1012 |
| D-limonene | 1032.09 | 1033 |
| eucalyptol | 1035.77 | 1033 |
| β-ocimene | 1039.84 | 1044 |
| trans-β-ocimene | 1050.53 | 1050 |
| acetophenone | 1071.3 | 1073 |
| *p*-cymene | 1089.01 | 1033 |
| unknown RT-13.13 | 1118.39 | ----- |
| unknown RT-13.87 | 1156.82 | ----- |
| unknown RT-15.15 | 1225.11 | ----- |
| unknown RT-15.31 | 1234.15 | ----- |
| thymol | 1250.78 | 1290 |
| *p*-cymen-7-ol | 1259.88 | 1278 |
| unknown RT-15.96 | 1270.26 | ----- |
| unknown RT-17.13 | 1337.57 | ----- |
| unknown RT-17.29 | 1346.91 | ----- |
| γ-elemene | 1402.27 | 1410 |
| β-guaiene | 1460.35 | 1479 |
| carotol | 1543.16 | 1594 |
| unknown RT-21.01 | 1583.39 | ----- |
| unknown RT-22.19 | 1692.64 | ----- |

**Supplementary Fig. S2:**
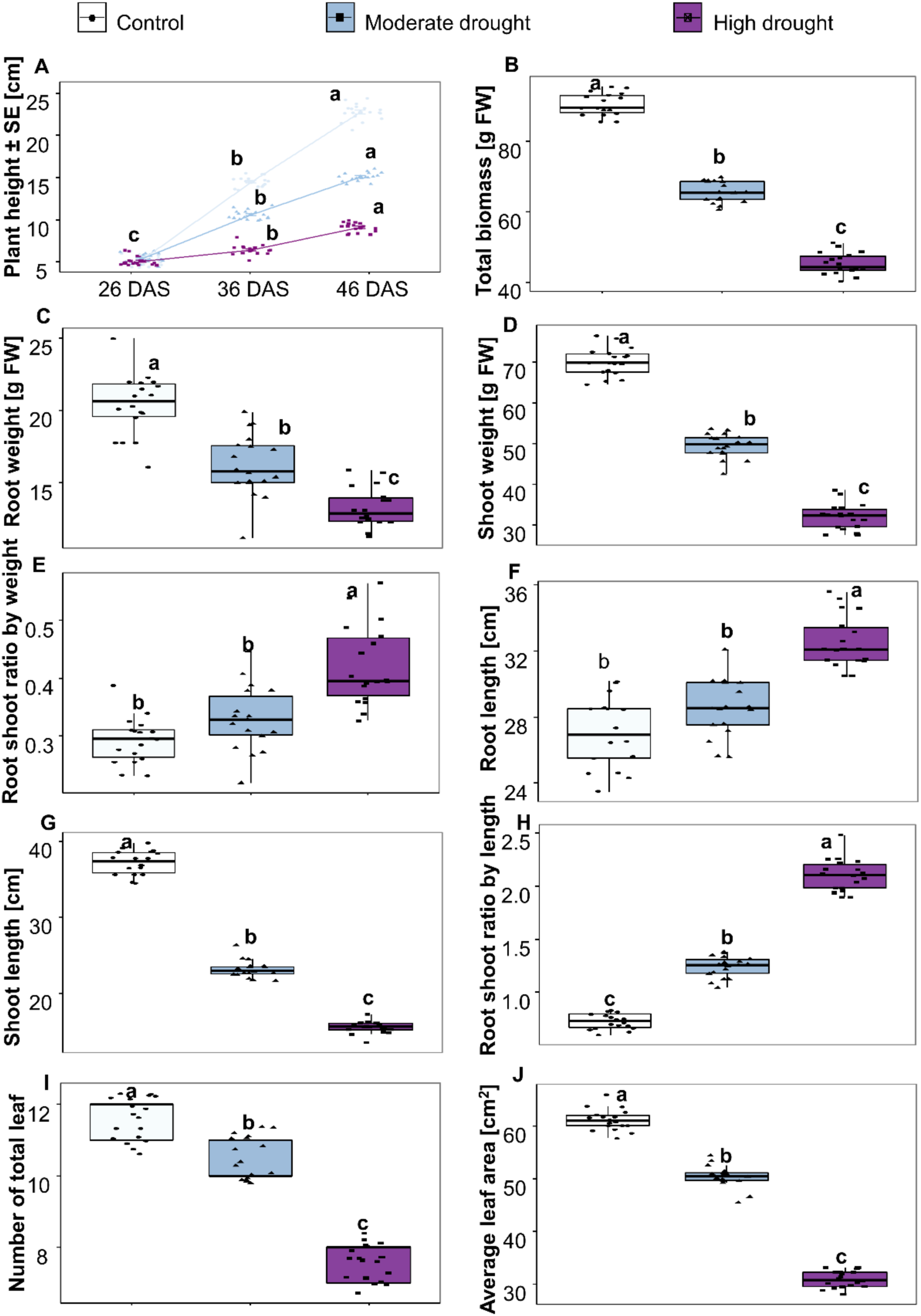
Effect of different magnitude of drought on sugarbeet plant morphological parameters. Drought treatments were implemented 27 days after sowing (DAS). Plants were harvested 58 DAS to record the parameters presented in B to J. Plant height (Test: GLMM) (A); Total biomass (Test: LMM) (B); Root weight (Test: LMM) (C); Shoot weight (Test: LMM) (D); Root shoot ratio by weight (Test: LMM) (E); Root length (Test: LMM) (F); Shoot length (Test: Kruskal-Wallis) (G); Root shoot ratio by length (Test: GLMM) (H); Number of leaves (Test: Kruskal-Wallis) (I); Average leaf area (Test: LMM) (J). Data points represent individual replicates, and different letters (*p* ≤ 0.05) indicate significance among treatments, n = 18

**Supplementary Fig. S3:**
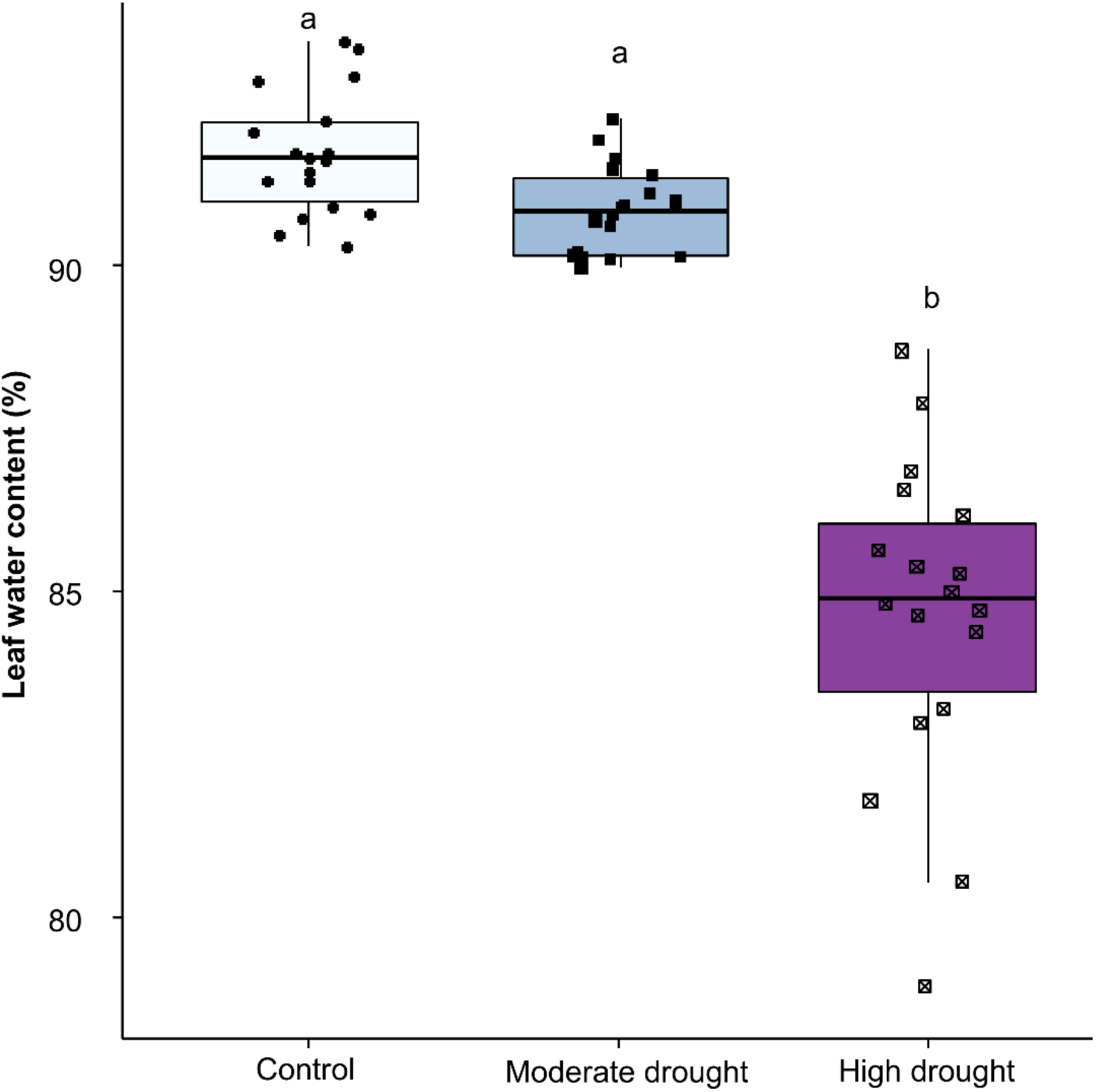
Effect of different magnitudes of drought on sugar beet leaf water content (Test: Kruskal-Wallis). Data points represent individual replicates, and different letters (*p* ≤ 0.05) indicate significance among treatments, n = 18.

**Supplementary Fig. S4:**
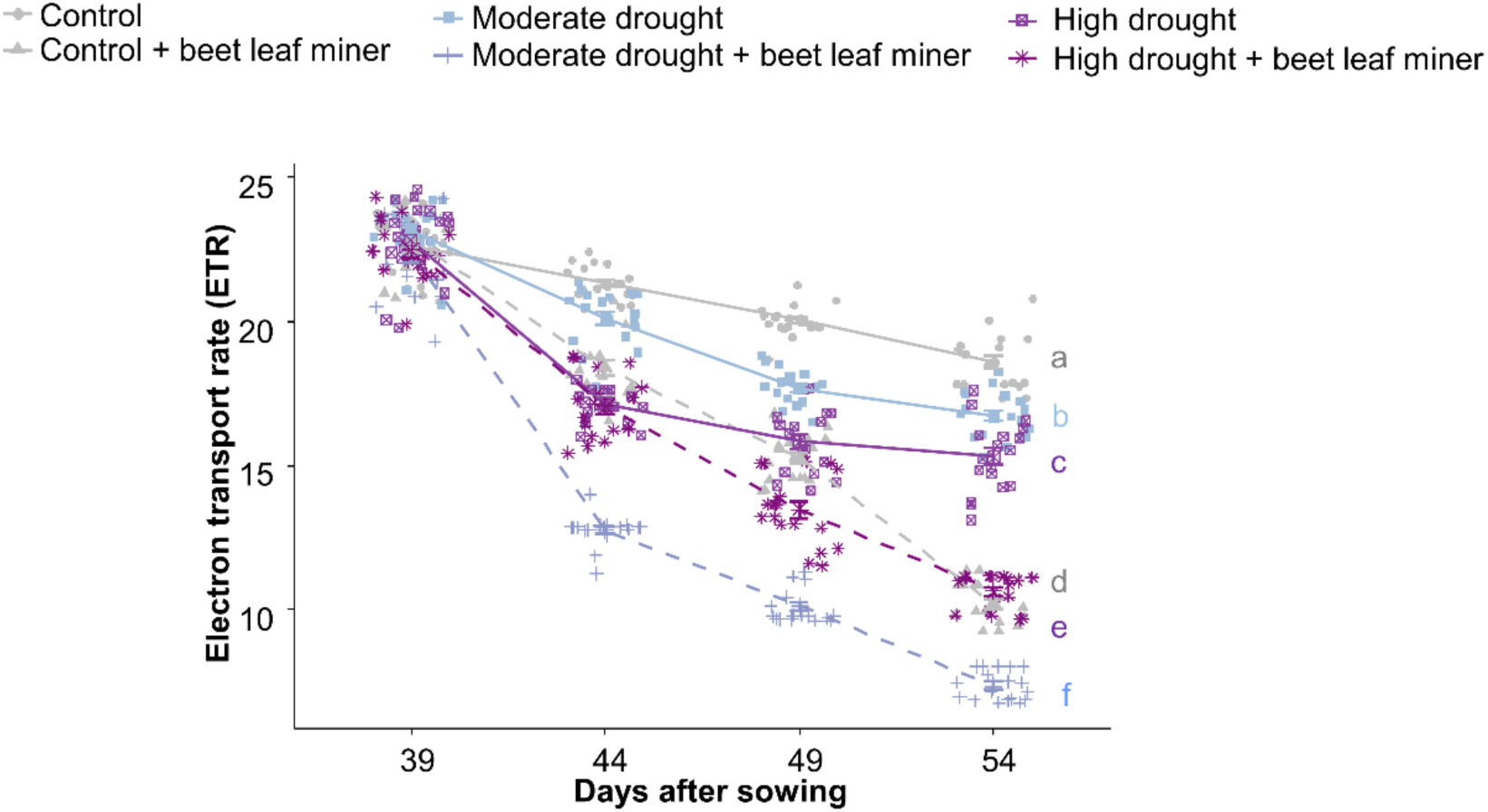
Effect of drought stress and beet miner infestation on sugar beet leaf photosynthetic efficiencies of electron transport rate (ETR) (Test: LM). Data point represents individual replicates, and different letters (*p* ≤ 0.05) indicate significance among treatments, n = 18.

**Supplementary Table S2:** Results of linear mixed models on leaf photosynthetic efficiencies parameters

| Photosynthetic efficiencies | Factors | Statistics |
| --- | --- | --- |
| Maximum quantum yield of PS II (Fv/Fm) | Drought | F2,408 = 601.66, p < 0.001 |
|  | Beet miner | F1,408 = 7656.06, p < 0.001 |
|  | Time | F3,408 = 4519.50, p < 0.001 |
|  | Drought × Beet miner | F2,408 = 898.51, p < 0.001 |
|  | Drought × Time | F6,408 = 126.87, p < 0.001 |
|  | Beet miner × Time | F3,408 = 1445.90, p < 0.001 |
|  | Drought × Beet miner × Time | F6,408 = 161.24, p < 0.001 |
| Quantum yield ($\Phi_{PSII}$) of PS II | Drought | F2,408 = 252.94, p < 0.001 |
|  | Beet miner | F1,408 = 2015.43, p < 0.001 |
|  | Time | F3,408 = 2069.20, p < 0.001 |
|  | Drought × Beet miner | F2,408 = 202.98, p < 0.001 |
|  | Drought × Time | F6,408 = 33.75, p < 0.001 |
|  | Beet miner × Time | F3,408 = 271.07, p < 0.001 |
|  | Drought × Beet miner × Time | F6,408 = 23.87, p < 0.001 |
| Electron transport rate (ETR) | Drought | F2,408 = 251.43, p < 0.001 |
|  | Beet miner | F1,408 = 1967.41, p < 0.001 |
|  | Time | F3,408 = 2031.44, p < 0.001 |
|  | Drought × Beet miner | F2,408 = 195.37, p < 0.001 |
|  | Drought × Time | F6,408 = 32.53, p < 0.001 |
|  | Beet miner × Time | F3,408 = 270.15, p < 0.001 |
|  | Drought × Beet miner × Time | F6,408 = 24.15, p < 0.001 |

**Supplementary Fig. S5:**
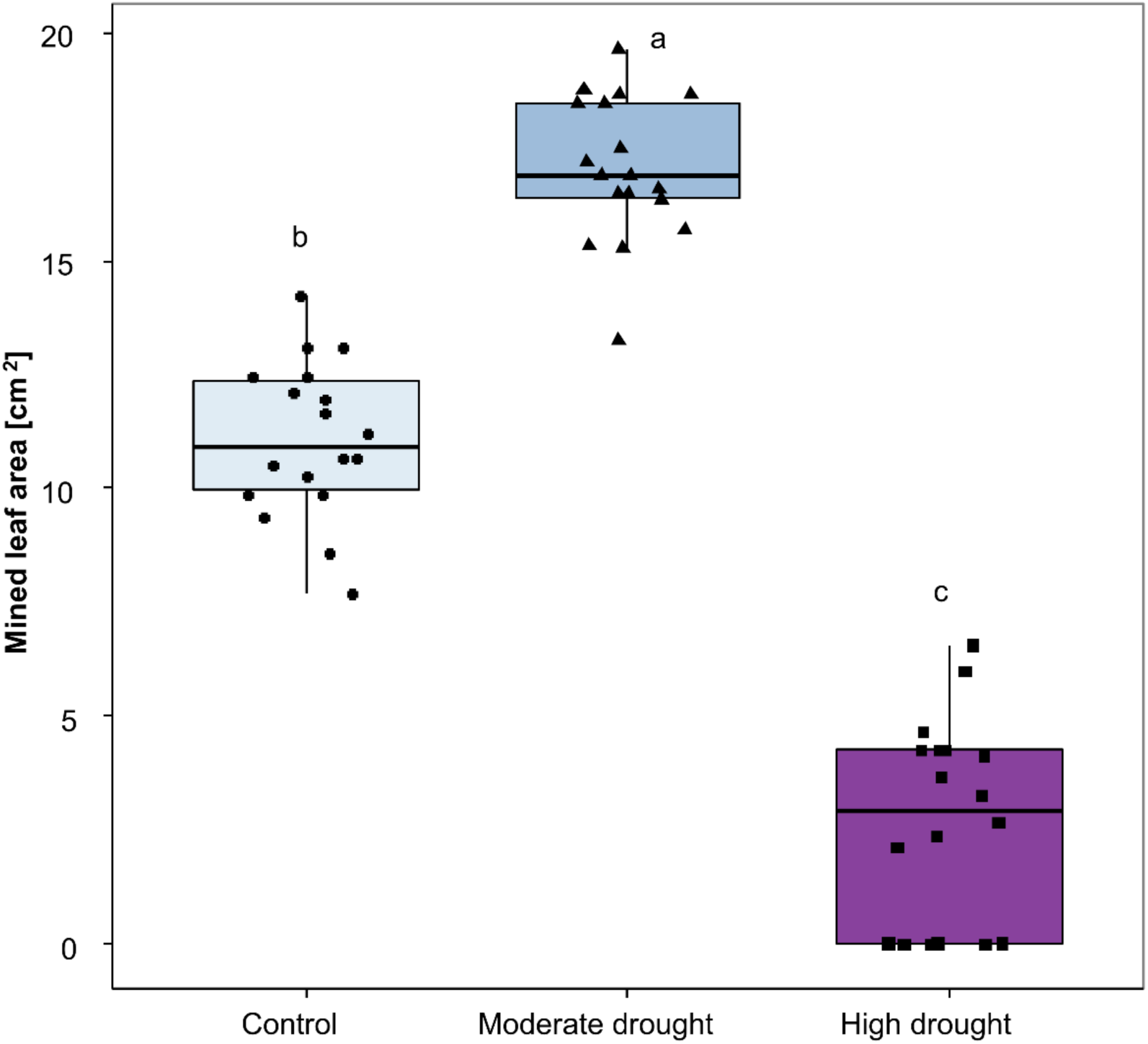
Effect of different magnitudes of drought on leaf area infestation by larvae of *Pegomya cunicularia* (Test: LMM). Data point represents individual replicates, and different letters (*p* ≤ 0.05) indicate significanct differences among treatments, n= 18.

**Supplementary Fig. S6:**
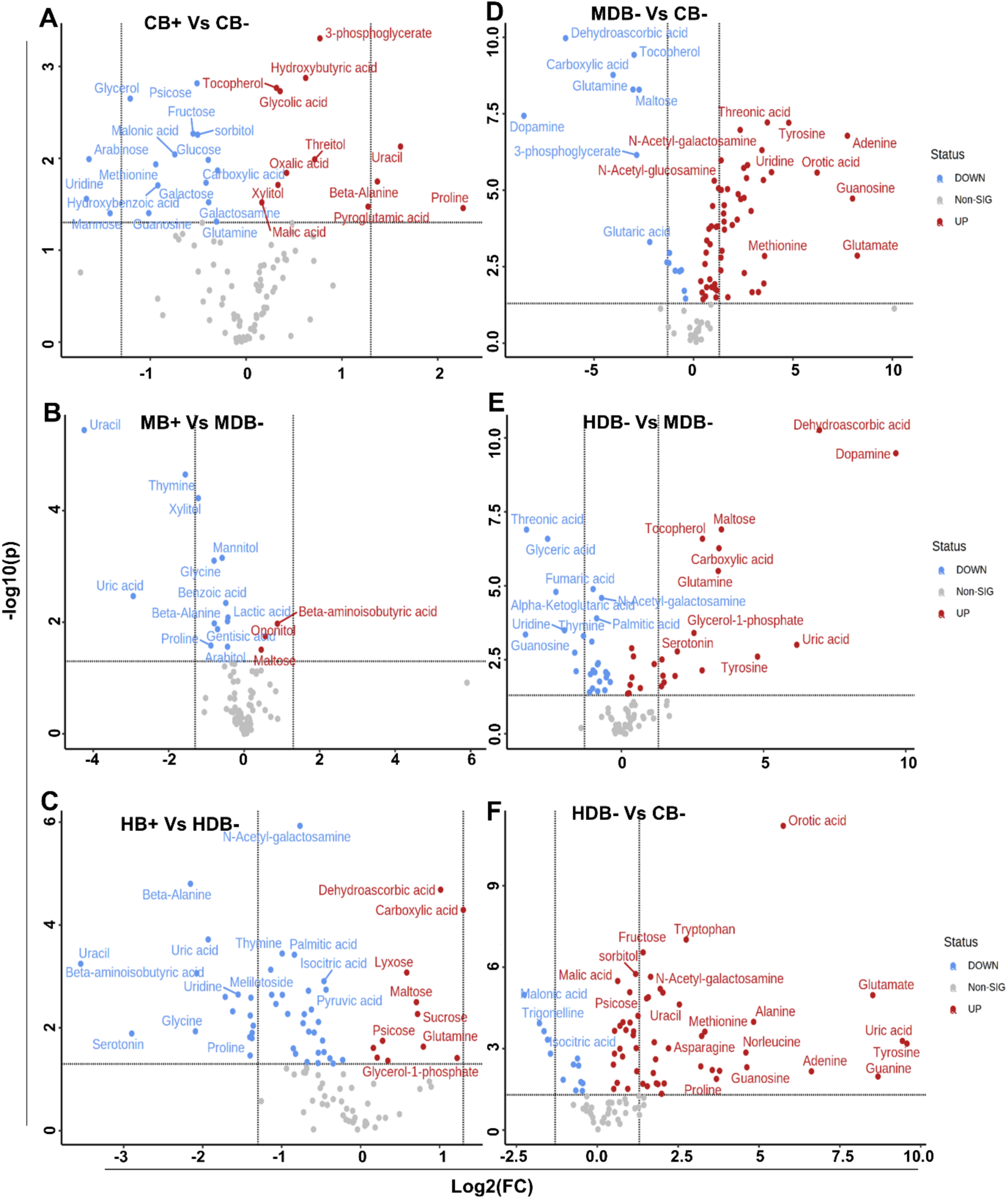
Effect of different magnitude of drought stress and beet miner infestation on central metabolites showing fold changes of up-regulated (in blue) and down-regulated (in red) metabolites. CB- = Control without beet leaf miner, CB+ = Control with beet leaf miner, MDB- = Moderate drought without beet leaf miner, MDB+ = Moderate drought with beet leaf miner, HDB- = High drought without beet leaf miner, HDB+ = High drought with beet leaf miner, n= 6.

**Supplementary Fig. S7:**
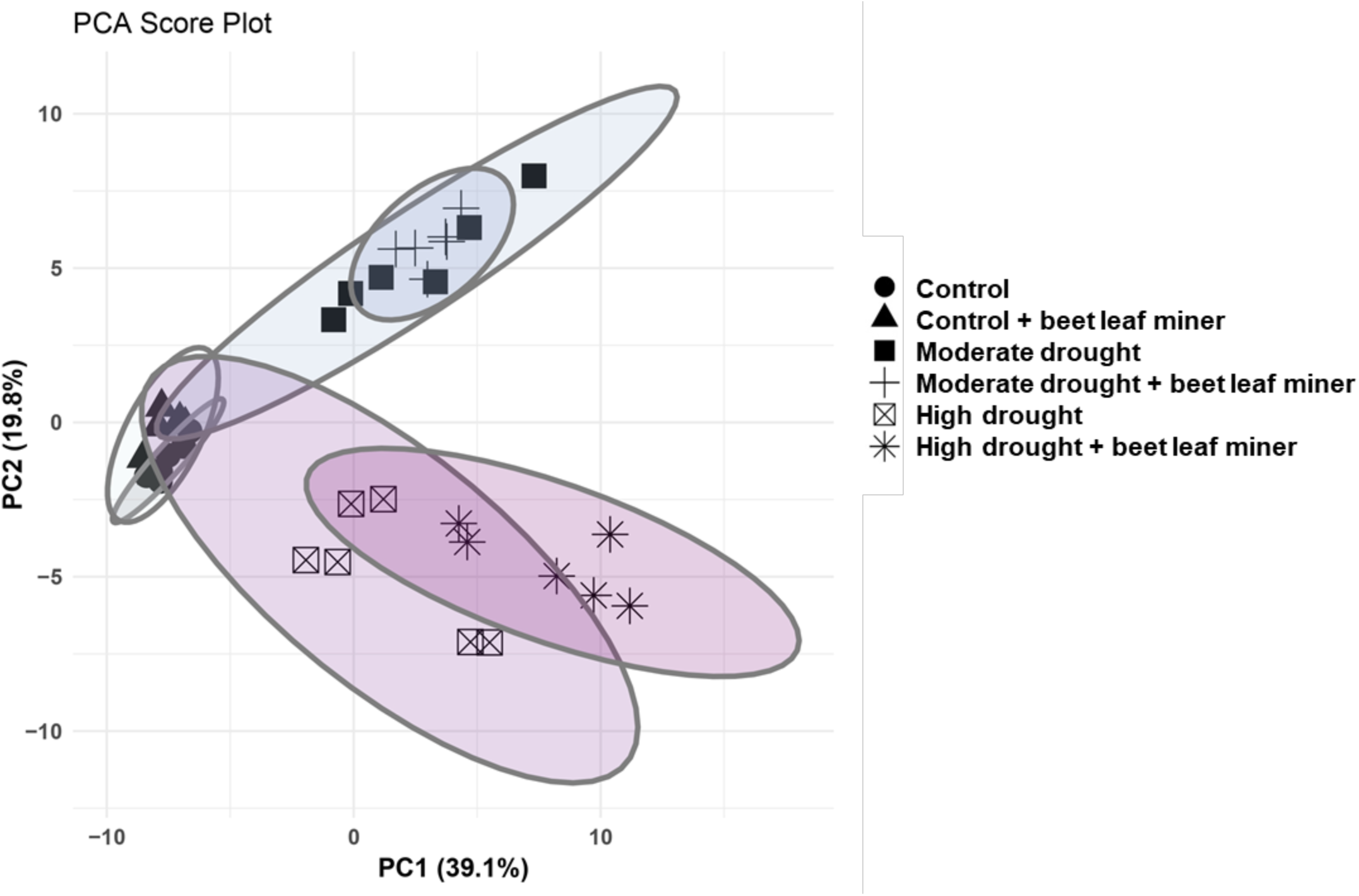
Principal component analysis (PCA) of central metabolites profiles of different treatments. Illustrated are the 95% confidence intervals for each group, n= 6.

**Supplementary Table S3:** Relative abundance of above-ground VOC emissions from whole sugar beet plants subjected to different treatments. Data is presented as relative abundance, calculated as the peak area ratio of the analyte to the internal standard (tetralin). CB- = Control without beet leaf miner, CB+ = Control with beet leaf miner, MDB- = Moderate drought without beet leaf miner, MDB+ = Moderate drought with beet leaf miner, HDB- = High drought without beet leaf miner, HDB+ = High drought with beet leaf miner. Differences among treatments were tested using Kruskal–Wallis test followed by Dunn’s post hoc test; different letters indicate significant differences among treatments (p ≤ 0.05, n = 6). χ² and p-values are shown in figure 5.

| Compound | VOC emission per plant<br>Relative abundance ± SE | | | | | |
| --- | --- | --- | --- | --- | --- | --- |
|  | CB- | CB+ | MDB- | MDB+ | HDB- | HDB+ |
| **trans-2-hexenal** | 0.08±0.005 abc | 0.32±0.21 ab | 0.03±0.008 c | 0.32±0.08 b | 0.09±0.01 abc | 0.07±0.02 ac |
| 4-hydroxy-4-methyl-2-pentanone | ----- | 0.01±0.006 | ----- | ----- | ----- | ----- |
| **3-hexenol** | 0.25±0.01 abc | 0.88±0.61 abc | 0.07±0.01 c | 0.91±0.14 a | 0.13±0.03 abc | 0.75±0.15 ac |
| α-pinene | 0.01±0.002 | 0.01±0.002 | ----- | 0.001±0.002 | ----- | ----- |
| **1-octen-3-ol** | 0.03±0.009 b | 0.10±0.02 a | ----- | ----- | ----- | ----- |
| **6-methyl-5-hepten-2-one** | 0.04±0.009 b | 0.14±0.03 a | ----- | ----- | ----- | ----- |
| β-myrcene | 0.01±0.003 | 0.04±0.01 | ----- | ----- | ----- | ----- |
| **octanal** | 0.01±0.001 ab | 0.04±0.01 a | ----- | 0.01±0.002 b | ----- | 0.02±0.002 a |
| **cis-3-hexenyl-1-acetate** | 3.97±0.64 ab | 4.14±0.13 abc | 0.38±0.02 c | 5.41±0.47 a | 0.79±0.17 ac | 4.28±0.05 abc |
| 3-carene | 0.06±0.01 | 0.15±0.05 | ----- | 0.11±0.01 | ----- | 0.08±0.004 |
| **D-limonene** | 0.05±0.008 ab | 0.09±0.01 b | 0.03±0.004ab | 0.02±0.0005 a | ----- | 0.04±0.02 ab |
| eucalyptol | ----- | 0.03±0.01 | ----- | ----- | ----- | ----- |
| **β-ocimene** | 0.08±0.02 ab | 0.27±0.07 a | ----- | 0.001±0.001 b | ----- | ----- |
| **trans-β-ocimene** | 0.04±0.01 b | 0.16±0.04 a | ----- | ----- | ----- | ----- |
| acetophenone | 0.02±0.005 | 0.06±0.01 | ----- | ----- | ----- | 0.06±0.01 |
| **p-cymene** | 0.01±0.001 b | 0.03±0.01 b | ----- | ----- | 0.05±0.008 ab | 0.10±0.01 a |
| unknown RT-13.13 | 0.04±0.01 | 2.16±1.3 | ----- | 0.18±0.07 | ----- | 0.22±0.07 |
| unknown RT-13.87 | ----- | 0.01±0.007 | ----- | ----- | ----- | ----- |
| unknown RT-15.15 | 0.03±0.01 | 0.05±0.01 | ----- | ----- | ----- | 0.04±0.01 |
| **unknown RT-15.31** | 0.08±0.01 ab | 0.18±0.06 ab | ----- | 0.03±0.004 b | ----- | 0.20±0.04 a |
| **thymol** | 0.13±0.03 ab | 0.43±0.12 a | 0.08±0.003ab | 0.05±0.02 b | 0.31±0.07 a | 0.21±0.06 a |
| **p-cymen-7-ol** | 0.36±0.05 ab | 0.68±0.16 a | 0.25±0.008ab | 0.19±0.03 b | 0.41±0.08 ab | 0.32±0.10 ab |
| **unknown RT-15.96** | 0.26±0.03 ab | 0.45±0.10 a | 0.17±0.007ab | 0.12±0.02 b | 0.28±0.04 ab | 0.22±0.06 ab |
| unknown RT-17.13 | ----- | 0.02±0.004 | ----- | ----- | ----- | ----- |
| unknown RT-17.29 | ----- | 0.02±0.006 | ----- | ----- | ----- | ----- |
| γ-elemene | ----- | 0.02±0.007 | ----- | 0.01±0.001 | ----- | ----- |
| β-guaiene | ----- | 0.07±0.1 | ----- | ----- | ----- | ----- |
| **carotol** | ----- | 0.04±0.01 a | ----- | 0.001±0.001 b | ----- | ----- |
| **unknown RT-21.01** | ----- | 0.82±0.43 a | ----- | 0.02±0.008 b | ----- | 0.02±0.006 b |
| **unknown RT-22.19** | 9.40±0.45 a | 11.76±0.77 a | 9.00±1.21 ab | 6.45±0.28 ab | 3.14±1.10 b | 2.56±0.97b |

**Supplementary Table S4:** Relative abundance of above-ground VOC emissions per gram of biomass from sugar beet plants subjected to different treatments. Data is presented as relative abundance, calculated as the peak area ratio of the analyte to the internal standard (tetralin). CB- = Control without beet miner, CB+ = Control with beet miner, MDB- = Moderate drought without beet miner, MDB+ = Moderate drought with beet miner, HDB- = High drought without beet leaf miner, HDB+ = High drought with beet leaf miner. Differences among treatments were tested using Kruskal–Wallis test followed by Dunn’s post hoc test; different letters indicate significant differences among treatments (p ≤ 0.05, n = 6). χ² and p-values are shown in figure 5.

| Compound | VOC emission standardized by plant biomass |  |  |  |  |  |
| --- | --- | --- | --- | --- | --- | --- |
| | Relative abundance ± SE | | | | | |
|  | CB- | CB+ | MDB- | MDB+ | HDB- | HDB+ |
| **trans-2-hexenal** | 0.01±0.0003 c | 0.02±0.01 ab | 0.01±0.001 c | 0.06±0.01 b | 0.02±0.004 ab | 0.02±0.006 ab |
| 4-hydroxy-4-methyl-2-pentanone | ----- | 0.001±0.0003 | ----- | ----- | ----- | ----- |
| **3-hexenol** | 0.02±0.001 bc | 0.05±0.03 abc | 0.01±0.002 c | 0.17±0.03 ab | 0.03±0.009abc | 0.22±0.05 a |
| α-pinene | 0.001±0.0001 | 0.001±0.0001 | ----- | 0.001±0.0003 | ----- | ----- |
| 1-octen-3-ol | 0.001±0.0007 | 0.01±0.001 | ----- | ----- | ----- | ----- |
| 6-methyl-5-hepten-2-one | 0.001±0.0007 | 0.01±0.001 | ----- | ----- | ----- | ----- |
| β-myrcene | 0.001±0.0002 | 0.001±0.0006 | ----- | ----- | ----- | ----- |
| **octanal** | 0.001±0.0001 a | 0.001±0.0005ab | ----- | 0.001±0.0003 a | ----- | 0.01±0.001 b |
| **cis-3-hexenyl-1-acetate** | 0.30±0.04 abc | 0.23±0.005 abc | 0.07±0.006 c | 1.01±0.11 ab | 0.20±0.05 bc | 1.27±0.10a |
| **3-carene** | 0.001±0.0008 c | 0.01±0.002 bc | ----- | 0.02±0.002 ab | ----- | 0.02±0.001 a |
| D-limonene | 0.001±0.0006 | 0.001±0.0008 | 0.01±0.001 | 0.001±0.0001 | ----- | 0.01±0.006 |
| eucalyptol | ----- | 0.00±0.0005 | ----- | ----- | ----- | ----- |
| **β-ocimene** | 0.01±0.002 a | 0.01±0.003 a | ----- | 0.001±0.0003 b | ----- | ----- |
| **trans-β-ocimene** | 0.001±0.001 a | 0.01±0.002 b | ----- | ----- | ----- | ----- |
| **acetophenone** | 0.001±0.0004 b | 0.001±0.0008ab | ----- | ----- | ----- | 0.02±0.005 a |
| **p-cymene** | 0.001±0.0001 c | 0.001±0.0005bc | ----- | ----- | 0.01±0.003 ab | 0.03±0.005 a |
| **unknown RT-13.13** | 0.001±0.0007 a | 0.11±0.06 ab | ----- | 0.03±0.01 ab | ----- | 0.07±0.02 b |
| unknown RT-13.87 | ----- | 0.001±0.0003 | ----- | ----- | ----- | ----- |
| unknown RT-15.15 | 0.001±0.0008 | 0.001±0.0007 | ----- | ----- | ----- | 0.01±0.003 |
| **unknown RT-15.31** | 0.01±0.001 b | 0.01±0.003 b | ----- | 0.01±0.0009 b | ----- | 0.06±0.006 a |
| **thymol** | 0.01±0.002 a | 0.02±0.006 ab | 0.01±0.001 ab | 0.01±0.003 a | 0.09±0.02 b | 0.07±0.02 ab |
| p-cymen-7-ol | 0.03±0.004 | 0.04±0.007 | 0.04±0.004 | 0.03±0.004 | 0.11±0.029 | 0.11±0.03 |
| **unknown RT-15.96** | 0.02±0.003 a | 0.02±0.004 ab | 0.03±0.002 ab | 0.02±0.003 ab | 0.07±0.01 b | 0.07±0.02 ab |
| unknown RT-17.13 | ----- | 0.001±0.0001 | ----- | ----- | ----- | ----- |
| unknown RT-17.29 | ----- | 0.001±0.0003 | ----- | ----- | ----- | ----- |
| γ-elemene | ----- | 0.001±0.0003 | ----- | 0.001±0.0003 | ----- | ----- |
| β-guaiene | ----- | 0.001±0.001 | ----- | ----- | ----- | ----- |
| carotol | ----- | 0.001±0.0007 | ----- | 0.001±0.0002 | ----- | ----- |
| unknown RT-21.01 | ----- | 0.04±0.02 | ----- | 0.001±0.001 | ----- | 0.001±0.001 |
| **unknown RT-22.19** | 0.70±0.04 bc | 0.65±0.03 c | 1.63±0.35 a | 1.19±0.02 ab | 0.84±0.31 abc | 0.71±0.20 c |

**Supplementary Fig. S8:**
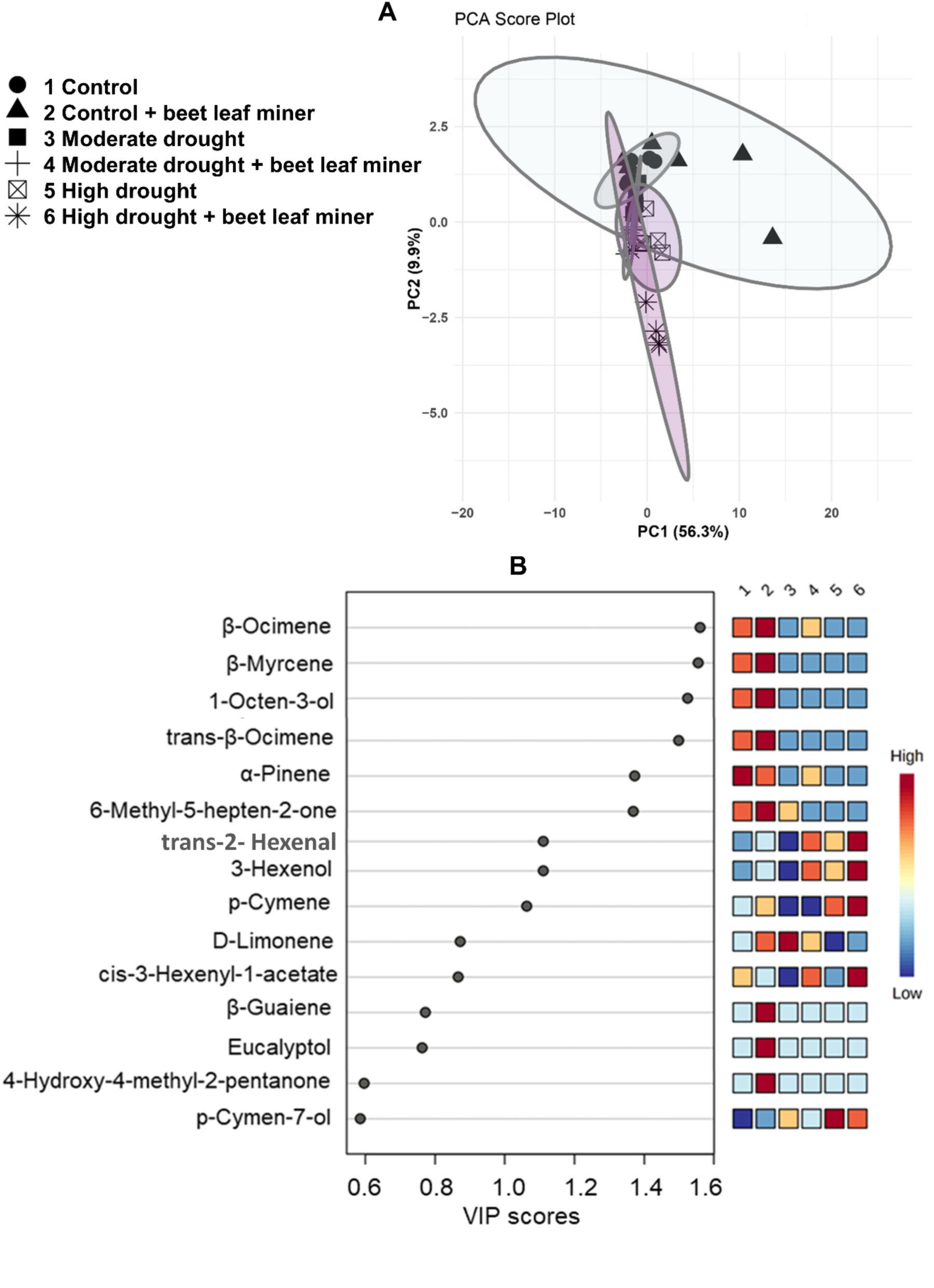
Score plot showing clustering of VOCs profiles in sugar beet in response to drought and beet miner stress for Principal component analysis (PCA) (A). Fifteen top VOCs are shown according to the variable importance of projection (VIP) score to different treatments (B). Colored boxes indicate the relative concentrations of the corresponding VOCs in each group. n= 6.

## References

Achhami BB, Reddy GVP, Hofland ML, Sherman JD, Peterson RKD, Weaver DK. 2021. Plant Volatiles and Oviposition Behavior in the Selection of Barley Cultivars by Wheat Stem Sawfly (Hymenoptera: Cephidae). Environmental Entomology 50, 940–947.

Agurla S, Gahir S, Munemasa S, Murata Y, Raghavendra AS. 2018. Mechanism of Stomatal Closure in Plants Exposed to Drought and Cold Stress. In: Iwaya-Inoue M, Sakurai M, Uemura M, eds. Advances in Experimental Medicine and Biology. Survival Strategies in Extreme Cold and Desiccation: Adaptation Mechanisms and Their Applications. Singapore: Springer, 215–232.

Anjum SA, Ashraf U, Tanveer M, et al. 2017. Drought Induced Changes in Growth, Osmolyte Accumulation and Antioxidant Metabolism of Three Maize Hybrids. Frontiers in Plant Science 8, 69.

Atkinson NJ, Urwin PE. 2012. The interaction of plant biotic and abiotic stresses: from genes to the field. Journal of Experimental Botany 63, 3523–3543.

Bezerra RHS, Sousa-Souto L, Santana AEG, Ambrogi BG. 2021. Indirect plant defenses: volatile organic compounds and extrafloral nectar. Arthropod-Plant Interactions 15, 467–489.

Bhattacharya A. 2021. Effect of Soil Water Deficit on Growth and Development of Plants: A Review. In: Bhattacharya A, ed. Soil Water Deficit and Physiological Issues in Plants. Singapore: Springer, 393–488.

Boergens E, Güntner A, Dobslaw H, Dahle C. 2020. Quantifying the Central European Droughts in 2018 and 2019 With GRACE Follow-On. Geophysical Research Letters 47, e2020GL087285.

Boggs CL, Freeman KD. 2005. Larval food limitation in butterflies: effects on adult resource allocation and fitness. Oecologia 144, 353–361.

Brilli F, Barta C, Fortunati A, Lerdau M, Loreto F, Centritto M. 2007. Response of isoprene emission and carbon metabolism to drought in white poplar (Populus alba) saplings. New Phytologist 175, 244–254.

Bruce TJA, Pickett JA. 2011. Perception of plant volatile blends by herbivorous insects – Finding the right mix. Phytochemistry 72, 1605–1611.

Bruce TJA, Wadhams LJ, Woodcock CM. 2005. Insect host location: a volatile situation. Trends in Plant Science 10, 269–274.

Carvajal Acosta AN, Agrawal AA, Mooney K. 2023. Plant water-use strategies as mediators of herbivore drought response: Ecophysiology, host plant quality and functional traits. Journal of Ecology 111, 687–700.

De Moraes CM, Mescher MC, Tumlinson JH. 2001. Caterpillar-induced nocturnal plant volatiles repel conspecific females. Nature 410, 577–580.

Deeba F, Pandey AK, Ranjan S, Mishra A, Singh R, Sharma YK, Shirke PA, Pandey V. 2012. Physiological and proteomic responses of cotton (Gossypium herbaceum L.) to drought stress. Plant Physiology and Biochemistry 53, 6–18.

Ebmeyer H, Fiedler-Wiechers K, Hoffmann CM. 2021. Drought tolerance of sugar beet – Evaluation of genotypic differences in yield potential and yield stability under varying environmental conditions. European Journal of Agronomy 125, 126262.

Erb M, Reymond P. 2019. Molecular Interactions Between Plants and Insect Herbivores. Annual Review of Plant Biology 70, 527–557.

Fàbregas N, Fernie AR. 2019. The metabolic response to drought. Journal of Experimental Botany 70, 1077–1085.

Fernández PC, Braccini CL, Dávila C, Barrozo RB, Aráoz MVC, Cerrillo T, Gershenzon J, Reichelt M, Zavala JA. 2019. The use of Leaf Surface Contact Cues During Oviposition Explains Field Preferences in the Willow Sawfly Nematus oligospilus. Scientific Reports 9, 4946.

Forieri I, Hildebrandt U, Rostás M. 2016. Salinity stress effects on direct and indirect defence metabolites in maize. Environmental and Experimental Botany 122, 68–77.

Franzke A, Reinhold K. 2011. Stressing food plants by altering water availability affects grasshopper performance. Ecosphere 2, 1–13.

Guerrieri E, Rasmann S. 2024. Insect herbivore induced above- and belowground plant communication: ecological and applied aspects. Entomologia Generalis, 1081–1090.

Hahn PG, Maron JL. 2018. Plant water stress and previous herbivore damage affect insect performance. Ecological Entomology 43, 47–54.

Hamann E, Blevins C, Franks SJ, Jameel MI, Anderson JT. 2021. Climate change alters plant–herbivore interactions. New Phytologist 229, 1894–1910.

Haslinger K, Mayer K. 2023. Early spring droughts in Central Europe: Indications for atmospheric and oceanic drivers. Atmospheric Science Letters 24, e1136.

Heil M. 2014. Herbivore-induced plant volatiles: targets, perception and unanswered questions. New Phytologist 204, 297–306.

Hillman S. 2022. Population Genetics and Ecology of the Sugar Beet Leaf Miners. doctoral, University of East Anglia. School of Biological Sciences.

Honda K. 1995. Chemical basis of differential oviposition by lepidopterous insects. Archives of Insect Biochemistry and Physiology 30, 1–23.

Igamberdiev AU, Kleczkowski LA. 2018. The Glycerate and Phosphorylated Pathways of Serine Synthesis in Plants: The Branches of Plant Glycolysis Linking Carbon and Nitrogen Metabolism. Frontiers in Plant Science 9.

Jaenike J. 1978. On optimal oviposition behavior in phytophagous insects. Theoretical Population Biology 14, 350–356.

Jones LC. 2022. Insects allocate eggs adaptively according to plant age, stress, disease or damage. Proceedings of the Royal Society B: Biological Sciences 289, 20220831.

Krasensky J, Jonak C. 2012. Drought, salt, and temperature stress-induced metabolic rearrangements and regulatory networks. Journal of Experimental Botany 63, 1593–1608.

Kuczyk J, Müller C, Fischer K. 2021. Plant-mediated indirect effects of climate change on an insect herbivore. Basic and Applied Ecology 53, 100–113.

Kulma A, Szopa J. 2007. Catecholamines are active compounds in plants. Plant Science 172, 433–440.

Kurepa J, Smalle JA. 2022. Auxin/Cytokinin Antagonistic Control of the Shoot/Root Growth Ratio and Its Relevance for Adaptation to Drought and Nutrient Deficiency Stresses. International Journal of Molecular Sciences 23, 1933.

Li Z-X, Tan J-F, Yao N, Xie R-H. 2024. From trade-off to synergy: how nutrient status modulates plant resistance to herbivorous insects? Advanced Biotechnology 2, 37.

Lin P-A, Chen Y, Ponce G, Acevedo FE, Lynch JP, Anderson CT, Ali JG, Felton GW. 2022. Stomata-mediated interactions between plants, herbivores, and the environment. Trends in Plant Science 27, 287–300.

Lin P-A, Kansman J, Chuang W-P, Robert C, Erb M, Felton GW. 2023. Water availability and plant–herbivore interactions. Journal of Experimental Botany 74, 2811–2828.

Loomis RS. 1997. On the utility of nitrogen in leaves. Proceedings of the National Academy of Sciences 94, 13378–13379.

Loreto F, Schnitzler J-P. 2010. Abiotic stresses and induced BVOCs. Trends in Plant Science 15, 154–166.

Maeda H, Dudareva N. 2012. The shikimate pathway and aromatic amino Acid biosynthesis in plants. Annual Review of Plant Biology 63, 73–105.

Marchin RM, Ossola A, Leishman MR, Ellsworth DS. 2020. A Simple Method for Simulating Drought Effects on Plants. Frontiers in Plant Science 10, 1715.

Mevi-Schütz J, Goverde M, Erhardt A. 2003. Effects of fertilization and elevated CO2 on larval food and butterfly nectar amino acid preference in Coenonympha pamphilus L. Behavioral Ecology and Sociobiology 54, 36–43.

Michelsen V. 1980. A revision of the beet leaf-miner complex, Pegomya hyoscyami s.lat. (Diptera: Anthomyiidae). Insect Systematics & Evolution 11, 297–309.

Obata T, Fernie AR. 2012. The use of metabolomics to dissect plant responses to abiotic stresses. Cellular and Molecular Life Sciences: CMLS 69, 3225–3243.

Pang, Z., Chong, J., Zhou, G., Morais D., Chang, L., Barrette, M., Gauthier, C., Jacques, PE., Li, S., and Xia, J. 2021. MetaboAnalyst 5.0: narrowing the gap between raw spectra and functional insights. Nucleic Acids Research, Pages W388–W396.

Pott DM, Osorio S, Vallarino JG. 2019. From Central to Specialized Metabolism: An Overview of Some Secondary Compounds Derived From the Primary Metabolism for Their Role in Conferring Nutritional and Organoleptic Characteristics to Fruit. Frontiers in Plant Science 10.

Rahman S, Rostás M, Vosteen I. 2025. Drought aggravates plant stress by favouring aphids and weakening indirect defense in a sugar beet tritrophic system. Journal of Pest Science 98, 549–564.

Ray RL, Ampim PAY, Gao M. 2020. Crop Protection Under Drought Stress. In: Jabran K, Florentine S, Chauhan BS, eds. Crop Protection Under Changing Climate. Cham: Springer International Publishing, 145–170.

Rocha M, Licausi F, Araújo WL, Nunes-Nesi A, Sodek L, Fernie AR, van Dongen JT. 2010. Glycolysis and the Tricarboxylic Acid Cycle Are Linked by Alanine Aminotransferase during Hypoxia Induced by Waterlogging of Lotus japonicus. Plant Physiology 152, 1501–1513.

Rojas JC, Virgen A, Cruz-López L. 2003. Chemical and Tactile Cues Influencing Oviposition of a Generalist Moth, Spodoptera frugiperda (Lepidoptera: Noctuidae). Environmental Entomology 32, 1386–1392.

Rusman Q, Cusumano A, Vosteen I. 2024. En route to resources: Foraging strategies of plant-associated insects to identify resources in complex dynamic environments. Functional Ecology 38, 1664–1682.

Saitta V, Rebora M, Piersanti S, Salerno G. 2024. Visual and chemical cues in the host plant selection of the melon ladybird Chnootriba elaterii (Coleoptera: Coccinellidae). Arthropod-Plant Interactions 18, 637–649.

Sander R. 2015. Compilation of Henry’s law constants (version 4.0) for water as solvent. Atmospheric Chemistry and Physics 15, 4399–4981.

Santiago-Salazar CM, Barrera JF, Rojas JC, Huerta-Palacios G, Escamilla-Prado E. 2022. Response of a specialist leaf miner insect to the environmental stress of its host plant. Arthropod-Plant Interactions 16, 329–339.

Sconiers WB, Eubanks MD. 2017. Not all droughts are created equal? The effects of stress severity on insect herbivore abundance. Arthropod-Plant Interactions 11, 45–60.

Sconiers WB, Rowland DL, Eubanks MD. 2020. Pulsed drought: The effects of varying water stress on plant physiology and predicting herbivore response. Crop Science 60, 2543–2561.

Sharma A, Shahzad B, Kumar V, Kohli SK, Sidhu GPS, Bali AS, Handa N, Kapoor D, Bhardwaj R, Zheng B. 2019. Phytohormones Regulate Accumulation of Osmolytes Under Abiotic Stress. Biomolecules 9, 285.

Shehzad M, Gulzar A, Staley JT, Tariq M. 2021. The effects of drought stress and type of fertiliser on generalist and specialist herbivores and their natural enemies. Annals of Applied Biology 178, 377–386.

Shin YK, Bhandari SR, Jo JS, Song JW, Lee JG. 2021. Effect of Drought Stress on Chlorophyll Fluorescence Parameters, Phytochemical Contents, and Antioxidant Activities in Lettuce Seedlings. Horticulturae 7, 238.

de Souza MWR, Ferreira EA, dos Santos JB, Soares MA, de Castro e Castro BM, Zanuncio JC. 2020. Fluorescence of chlorophyll a in transgenic maize with herbicide application and attacked by Spodoptera frugiperda (Lepidoptera: Noctuidae). Phytoparasitica 48, 567–573.

Szabó K, Zubay P, Németh-Zámboriné É. 2020. What shapes our knowledge of the relationship between water deficiency stress and plant volatiles? Acta Physiologiae Plantarum 42, 130.

Tariq M, Wright DJ, Bruce TJA, Staley JT. 2013. Drought and Root Herbivory Interact to Alter the Response of Above-Ground Parasitoids to Aphid Infested Plants and Associated Plant Volatile Signals. PLOS ONE 8, e69013.

Turlings TCJ, Davison AC, TamÒ C. 2004. A six-arm olfactometer permitting simultaneous observation of insect attraction and odour trapping. Physiological Entomology 29, 45–55.

Viric Gasparic H, Lemic D, Drmic Z, Cacija M, Bazok R. 2021. The Efficacy of Seed Treatments on Major Sugar Beet Pests: Possible Consequences of the Recent Neonicotinoid Ban. Agronomy 11, 1277.

Wei J, Zou L, Kuang R, He L. 2000. Influence of Leaf Tissue Structure on Host Feeding Selection by Pea Leafminer Liriomyza huidobrensis (Diptera: Agromyzidae). Zoological Studies 39, 295–300.

Weldegergis BT, Zhu F, Poelman EH, Dicke M. 2015. Drought stress affects plant metabolites and herbivore preference but not host location by its parasitoids. Oecologia 177, 701–713.

Wittstock U, Gershenzon J. 2002. Constitutive plant toxins and their role in defense against herbivores and pathogens. Current Opinion in Plant Biology 5, 300–307.

Yang X, Lu M, Wang Y, Wang Y, Liu Z, Chen S. 2021. Response Mechanism of Plants to Drought Stress. Horticulturae 7, 50.

Yao J, Sun D, Cen H, Xu H, Weng H, Yuan F, He Y. 2018. Phenotyping of Arabidopsis Drought Stress Response Using Kinetic Chlorophyll Fluorescence and Multicolor Fluorescence Imaging. Frontiers in Plant Science 9.

Zebelo SA, Maffei ME. 2015. Role of early signalling events in plant–insect interactions. Journal of Experimental Botany 66, 435–448.

Zhuang J, Wang Y, Chi Y, Zhou L, Chen J, Zhou W, Song J, Zhao N, Ding J. 2020. Drought stress strengthens the link between chlorophyll fluorescence parameters and photosynthetic traits. PeerJ 8, e10046.

## References

Brooks ME, Kristensen K, Benthem KJ van, et al (2017) glmmTMB Balances Speed and Flexibility Among Packages for Zero-inflated Generalized Linear Mixed Modeling. The R Journal 9:378–400

Florian Hartig (2022) DHARMa: Residual Diagnostics for Hierarchical (Multi-Level / Mixed) Regression Models

Hothorn T, Bretz F, Westfall P (2008) Simultaneous Inference in General Parametric Models. Biom J 50:346–363. 10.1002/bimj.200810425

José Pinheiro JP, Douglas Bates, Saikat DebRoy, et al (2023) nlme: Linear and Nonlinear Mixed Effects Models

Lenth RV, Bolker B, Buerkner P, et al (2023) emmeans: Estimated Marginal Means, aka Least-Squares Means

Mangiafico S (2023) rcompanion: Functions to Support Extension Education Program Evaluation

